# Membrane cholesterol interferes with tyrosine phosphorylation but facilitates the clustering and signal transduction of EGFR

**DOI:** 10.1101/2021.08.28.457965

**Authors:** Michio Hiroshima, Mitsuhiro Abe, Nario Tomishige, Françoise Hullin-Matsuda, Asami Makino, Masahiro Ueda, Toshihide Kobayashi, Yasushi Sako

## Abstract

Epidermal growth factor receptor (EGFR) activates major cell signaling pathways that regulate various cell responses. Its dimerization and clustering coupled with its lateral mobility are critical for EGFR function, but the contribution of the plasma membrane environment to EGFR function is unknown. Here we show, using single-molecule analysis, that EGFR mobility and clustering are altered by the depletion of cholesterol or sphingomyelin, major lipids of membrane subdomains, causing significant changes in EGFR signaling. When cholesterol was depleted, the subdomain boundary in EGFR diffusion disappeared, the fraction of EGFR pre-dimers was increased, and the ligand-induced phosphorylation of EGFR was enhanced. In addition, the depletion of either lipid prevented the formation of immobile clusters after EGF association and decreased the phosphorylation of downstream proteins. Our results revealed that cholesterol plays dichotomous roles in the signaling pathway of EGFR and that clustering in the membrane subdomains is critical for EGFR signal transduction.

## Introduction

Epidermal growth factor receptor (EGFR), a receptor tyrosine kinase, is a major regulator of several intracellular signaling cascades by receiving extracellular ligands at the plasma membrane. EGFR signaling transfers information to the well-known RAS-MAPK pathway, inducing essential cell responses such as proliferation, differentiation, migration, apoptosis, and others. Defects in EGFR function affect cellular responses, often inducing hyperactivated signaling to causes carcinoma and other diseases (Carpenter et al., 1978; Lemmon and Schlessinger, 2010). Ligand-bound EGFR is auto-phosphorylated when EGFR takes a dimer structure to activate downstream signaling. Even before the ligand binding, dimer formation (pre-dimer) occurs (Hiroshima et al., 2012, 2018; Martin-Fernandez et al., 2002; Teramura et al., 2006; Yu et al., 2002), but usually the dimer cannot evoke whole cell activation and instead results in only higher ligand-affinity than the monomer. The ligand binding alters the pre-dimer state to an active dimer state capable of auto-phosphorylation. Previous studies have also indicated the existence of receptor pre-clusters larger than the dimer (Tao and Maruyama, 2008; Webb et al., 2008). Since the dimerization and cluster formation are driven by a molecular collision during diffusion along the plasma membrane, the membrane environment is reflected in the EGFR behavior (Arkhipov et al., 2013; Lin et al., 2016; Valley et al., 2015) and affects the EGFR signaling (Lajoie et al., 2007). To elucidate the membrane environment effects, observation of the spatiotemporal behaviors of individual EGFR molecules in cells is necessary.

Single-molecule trajectory analysis combined with the hidden Markov model (HMM) based on machine learning methods (Okamoto and Sako, 2012; Rabiner, 1989) has been used to infer molecular state transitions along the trajectory of an individual molecule (Chung et al., 2010; Low-Nam et al., 2011; Persson et al., 2013). EGFR was shown to transit between three motional states; namely, immobile, slow-, and fast-mobile, which were determined in terms of the size of the diffusion coefficient (Hiroshima et al., 2018; Yasui et al., 2018). We previously found that EGFR molecules in the slow-mobile state showed a confined diffusion surrounding the trajectory of the immobile state in which the position of the molecules fluctuated within a confined area (~60 nm) only two times larger than the localization accuracy. Fast-mobile EGFR molecules moved in space between the slow-mobile compartment with simple diffusion. The clustering states, which correspond to the number of EGFR molecules moving together, were determined from the brightness of the GFP fluorescence probing EGFR. After EGF stimulation, the EGFR clustering state shifted initially from monomer to dimer and subsequently to larger clusters concurrent with a shift to a slower mobility state. Immobile clusters are the primary interaction sites with the downstream protein GRB2. The dissociation kinetics between EGFR and GRB2 is specifically slower in immobile clusters, suggesting they play a significant role in the signal transduction in cells (Hiroshima et al., 2018). Overall, these results suggested that lateral mobility, clustering, and signal activation are closely correlated in EGFR.

The confinement size of the slow-mobile diffusion we have observed (~200 nm) is equivalent to the size often reported for lipid rafts (a membrane subdomain), which are a liquid-ordered phase segregated from the bulk region (liquid disordered phase) of the plasma membrane (Semrau and Schmidt, 2009). However, little is understood about the relationship between EGFR behavior and the membrane environment. Cholesterol and sphingomyelin are well-known major components of lipid rafts and can be depleted from the plasma membrane by treatment with methyl-β-cyclodextrin (MβCD), which extracts cholesterol from the membrane to its hydrophobic cavity (Zidovetzki and Levitan, 2007), or sphingomyelinase (SMase), which catalyzes the breakdown of sphingomyelin to phosphorylcholine and ceramide; ceramide is then converted to sphingosine and sphingosine-1-phosphate (S1P) and transported out of the cells (Hannun and Obeid, 2008; Sasset et al., 2016). MβCD and sphingomyelinase treatments have been reported to disperse and affect the physical properties of lipid rafts, respectively (Cremesti et al., 2002; Smith et al., 2010), indicating they alter EGFR behavior. These perturbations offer information on how cellular signaling is affected by the plasma membrane environment through the dynamics of EGFR behavior at the molecular level.

To understand the dependency of EGFR behavior, including the localization, mobility, clustering, and their coupling, on the membrane structure, the present study employed single-molecule analysis while depleting cholesterol or sphingomyelin. Furthermore, assessments of the receptor and its downstream activity were carried out to reveal the correlation between EGFR behavior and cellular signaling.

## Results

### Cholesterol-but not sphingomyelin-depletion enhanced EGF-induced EGFR phosphorylation

CHO-K1 cells were transfected with EGFR-GFP and treated with MβCD or sphingomyelinase for the lipid depletion. Cholesterol was reduced to 33% and 16% with 5 and 10 mM MβCD treatment, respectively, according to GC-FID or GC-MS measurements (Fig. 1a). Similar depletion was also confirmed by the exogenous addition of fluorescent EGFP-labeled θ toxin, a probe of free cholesterol (Fig. 1b), that binds to the cells. Sphingomyelin was reduced to 18% by sphingomyelinase treatment (Fig. 1b) based on the fluorescence of GFP-labeled lysenin, a specific probe of sphingomyelin (Fig. 1b). The observed fluorescence of θ toxin-GFP in sphingomyelinase-treated cells and lysenin-GFP in MβCD-treated cells were the same as in non-treated cells (Fig. S1), indicating that the specific depletion of one lipid neither affected the content of the other lipid in cells.

**Fig. 1.**
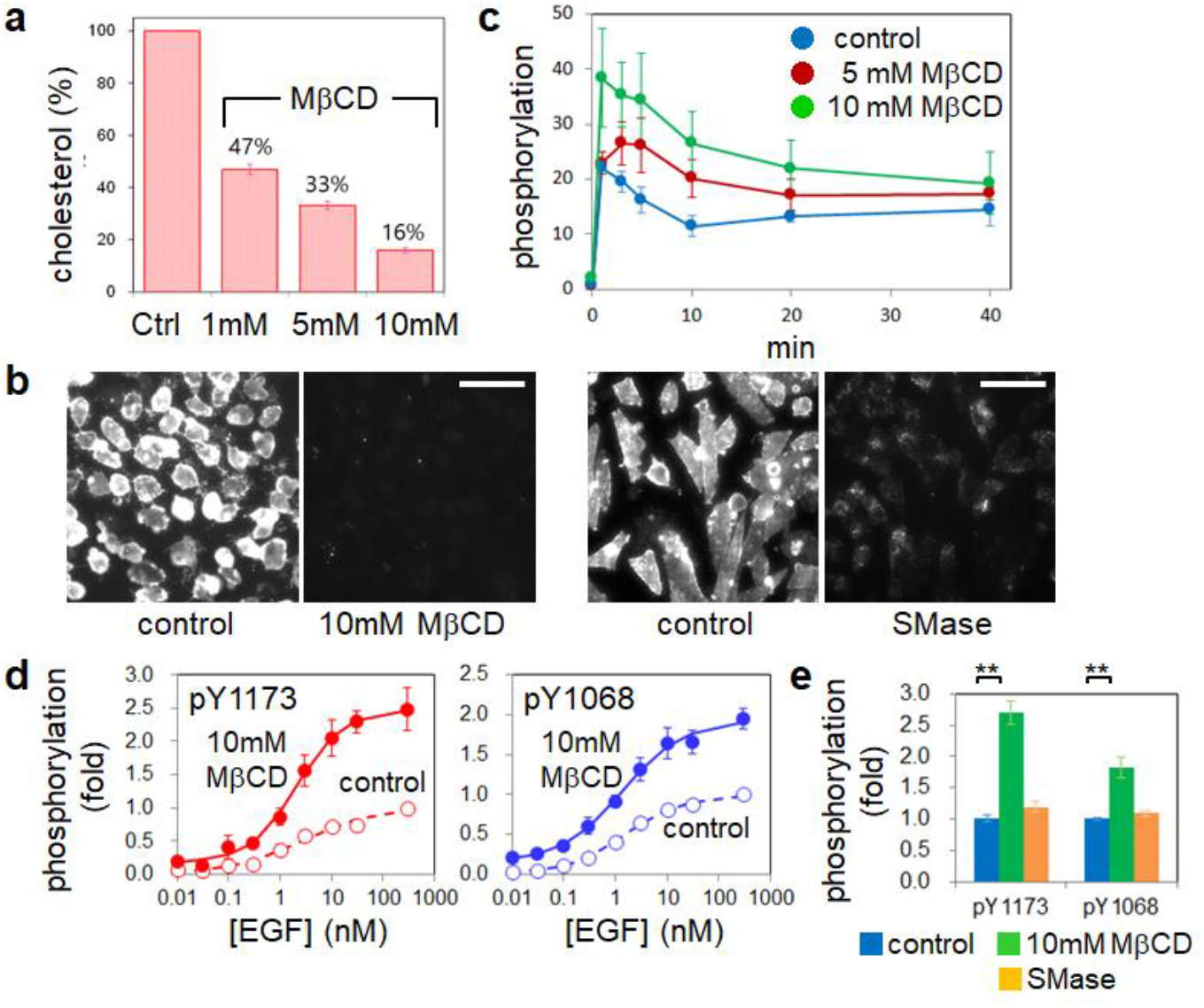
Lipid-depletion and EGFR phosphorylation. **a**. Cholesterol content in MβCD-treated cells. The amount of cholesterol was normalized to that in non-treated cells. **b**. Fluorescence images of wild-type CHO-K1 cells labeled with GFP-conjugated lipid probes. Left: θ toxin-labeled cells. Right: Lysenin-labeled cells. Scale bars, 50 μm. **c**. Time-course of the EGF-induced EGFR phosphorylation (pY1173) (n = 3 trials). Fold-changes relative to phosphorylation at 0 min are indicated. **d**. Dose-response curves for EGF-induced tyrosine phosphorylation in EGFR. **e**. Comparison between the phosphorylation levels at 2 min after 30 nM EGF stimulation. ** p < 0.01 (t-test). **a-e**, Error bars: SE. All data points are shown in Fig. S2.

The time-course of EGFR phosphorylated at Y1173 in cholesterol-depleted and control cells reached a maximum at 1-3 minutes after EGF stimulation according to Western blotting results (Fig. 1c and S2a). The phosphorylation level under cholesterol-depletion was higher than that in the control condition and was dependent on the MβCD concentration. The phosphorylation of EGFR at Y1173 and Y1068 was increased by the cholesterol depletion two minutes after the stimulation (Fig. 1d). The half-maximal effective concentrations (EC_50_) of EGF was almost the same between control and cholesterol-depleted conditions: 1.9 nM and 2.1 nM for pY1173, and 1.5 nM and 1.3 nM for pY1068, respectively. Hill coefficients indicating no cooperativity (0.6-1.0) were not changed by the depletion. After 30 nM EGF stimulation, the cholesterol-depletion condition induced 1.8-fold and 2.7-fold higher phosphorylation of Y1068 and Y1173, respectively, compared with the control condition (Fig.1e and S2b). Sphingomyelin depletion did not affect the phosphorylation of Y1068 or Y1173 significantly (Fig. 1e).

### Cholesterol-depletion enlarged the slow-mobile region before EGF stimulation

To understand how the differences in phosphorylation arose among the control and cholesterol-depleted conditions with consideration of EGFR behavior, we applied single-molecule imaging of EGFR-EGFP on the plasma membrane of living cells. We analyzed the trajectories of individual fluorescent spots using an HMM-based machine learning method to assign every step along the trajectories with specific motional and clustering states. The movements consisted of immobile, slow-, and fast-mobile states in all lipid conditions (Fig. S3 and Table S1). The depletion of either lipid increased the diffusion coefficient in the immobile state and the fraction of the slow-mobile state while decreasing the fraction of the fast-mobile state. Sphingomyelin depletion also increased the diffusion coefficient of the fast-mobile state. The observed changes in the diffusion coefficients were consistent with previous reports indicating that membrane fluidity is reduced by the addition of cholesterol (Tabas, 2002) or sphingomyelin (Makdissy et al., 2015). Trajectories of the lateral motion (Fig. 2) showed that cholesterol-depletion enlarged the diffusion region especially during the slow-mobile state (Fig. 2a, orange) but that the sphingomyelin-depletion had little effect. These trajectories reflect the properties of the time evolution of the mean square displacement (MSD; Fig. 2b and S4).

**Fig. 2.**
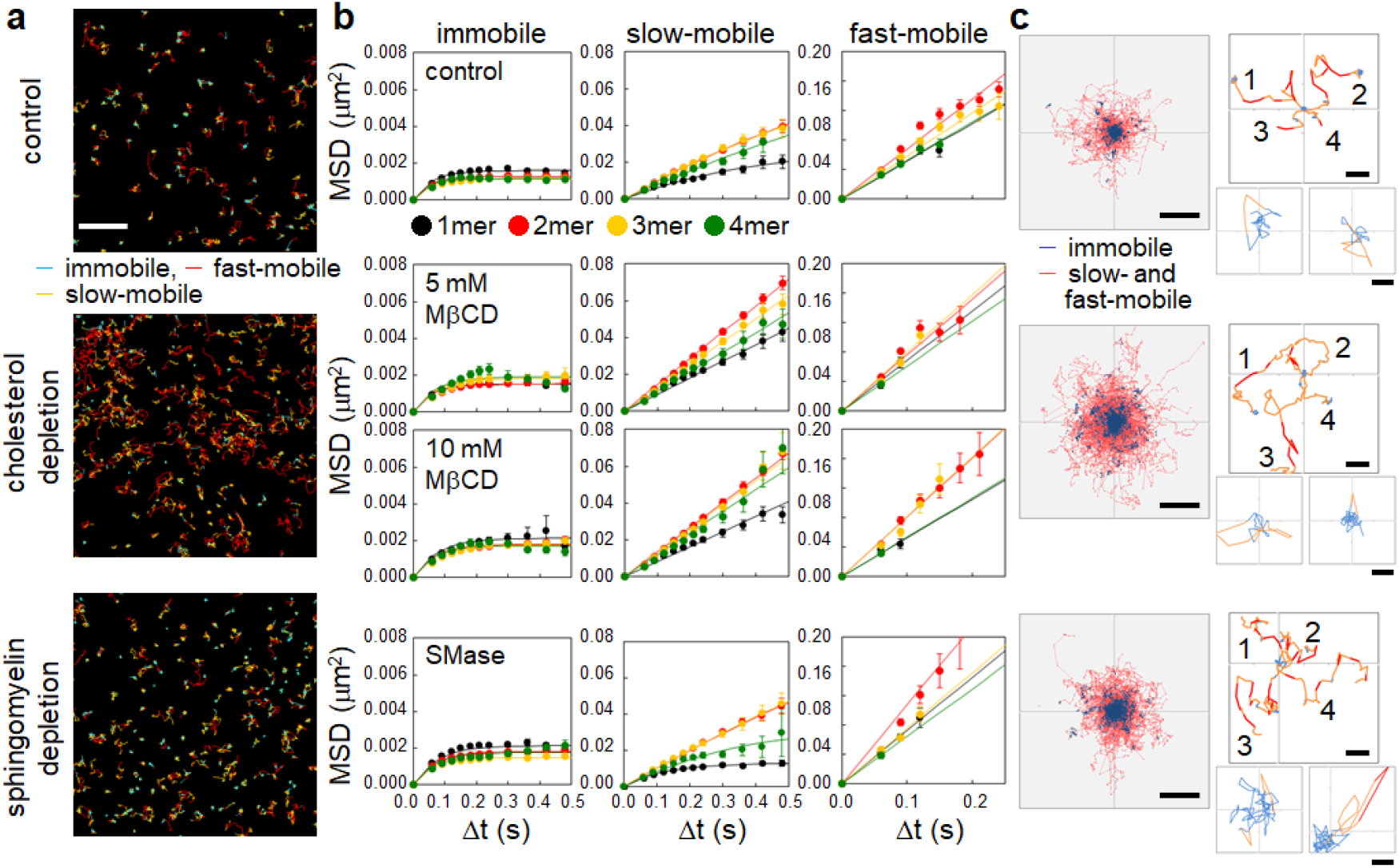
Lipid-depletion and EGFR mobility in the absence of EGF. **a**. Trajectories of EGFR particles in three state transitions. Scale bar, 2 μm. **b**. MSD-Δt plots for each motional and clustering state. Data are shown as the mean and SE. All single-cell data points are shown in Fig. S4. **c**. Trajectories of 500 particles up to 50 frames (1.5 sec) originated from the immobile region were superimposed (left). Scale bar, 500 nm. Typical trajectories were extracted in the right panels. The number denotes each trajectory. Scale bars, 300 nm. In the smaller panels, trajectories that came back to the identical immobile region are shown. Scale bars, 50 nm.

MSD profiles of confined and simple diffusion were fitted with the following equations, respectively,

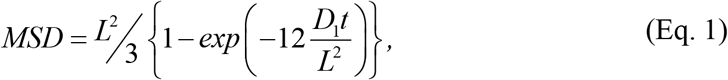

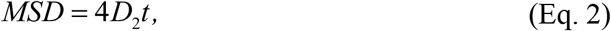

where *D*_1_ and *D*_2_ are the diffusion coefficients, *L* is the confinement length, and *t* is the diffusion time. The suitable diffusion mode for each MSD profile was determined using Akaike’s information criterion (AIC), which was calculated using the equation below,

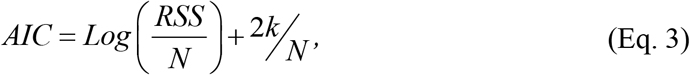

where *RSS* is the residual sum of squares between the data and the model, *N* is the number of data points, and *k* is the number of parameters. The model with the higher AIC was selected. In the control condition, confined diffusion was observed in the immobile and slow-mobile states, but the fast-mobile state showed simple diffusion. The confinement lengths (*L*) of the immobile state were 60 nm for all cluster sizes (monomer, dimer, and higher-order clusters). *L* for the slow-mobile state was equivalent to the size of well-known membrane subdomains (including lipid rafts). Finally, the mobility of the monomer (~310 nm) was more confined than the mobility of the other clusters (~570 nm).

When cholesterol was depleted, the diffusion mode in the slow-mobile state was altered from confined to simple diffusion for all cluster sizes. The cholesterol depletion had little effect on the MSD of both the immobile and fast-mobile states or on the distance between the centers of the immobile state regions (Fig. S5). When sphingomyelin was depleted, the slow mobile EGFR still showed confined diffusion, but the confinement was less in comparison with the control condition (~720 nm for ≥ dimer). In Fig. 2c, individual trajectories were superimposed to exhibit the expanding diffusion area of EGFR, for which the center of the first immobile state in each trajectory was shifted to the origin. The second and later immobile states are seen as small islands separated from the first region, and the slow- and fast-mobile states were distributed around the immobile states (Fig. 2c, left). Typical trajectories (Fig. 2c, right) rarely showed a direct transition between the immobile and fast-mobile states, as observed in the transition probability (Table S1). Transitions between the immobile and slow-mobile states often occurred at the surrounding boundary of the region for an immobile state, suggesting that an EGFR particle in the immobile state was trapped within a 60-nm membrane subdomain that was stable during the observation time. Boundaries between the slow-mobile and fast-mobile regions were obscure. Thus, the small core area surrounded by a region several hundred nanometers wide forms a membrane subdomain in which EGFR molecules are confined. In the space between the subdomains, EGFR molecules are allowed to diffuse in the fast-mobile state. The cholesterol depletion loosens the confinement and enlarges the range of slow-mobile motion (Fig. 2c, left middle), possibly leading to occasional subdomain fusion. In the case of sphingomyelin-depletion,slight expansion of the diffusion area was observed (Fig. 2c, left bottom), reflecting the increase in both the confinement length of the slow-mobile state and the diffusion coefficient of the fast-mobile state.

### Cholesterol depletion increased the slow-mobile EGFR pre-dimer

The cluster size distribution, which was obtained from the HMM analysis, showed that EGFR dimer and higher-order clusters existed even without ligand stimulation and thus could be called pre-dimer and pre-clusters, respectively. The pre-dimer is responsible for increasing the sensitivity of EGF signaling (Hiroshima et al., 2012; Teramura et al., 2006), but spontaneous auto-phosphorylation hardly occurs. Ligand binding alters the pre-dimer to an active configuration (Hofman et al., 2010), enabling auto-phosphorylation. We found that when cholesterol was depleted, the fraction of pre-dimer in the slow-mobile state was significantly increased (1.4-fold), but the fractions of monomers and higher-order clusters (≥ trimer) were unchanged (Fig. 3a and S6), suggesting an upshift in dimerization affinity between EGFR monomers and the destabilization of clusters to the dimer. The fractions from monomer to tetramer were reduced in the fast-mobile state, but no change was observed in the immobile state (Table S2). These changes increased the total slow-mobile fraction 1.3-fold (Fig. S3b and Table S1).

**Fig. 3.**
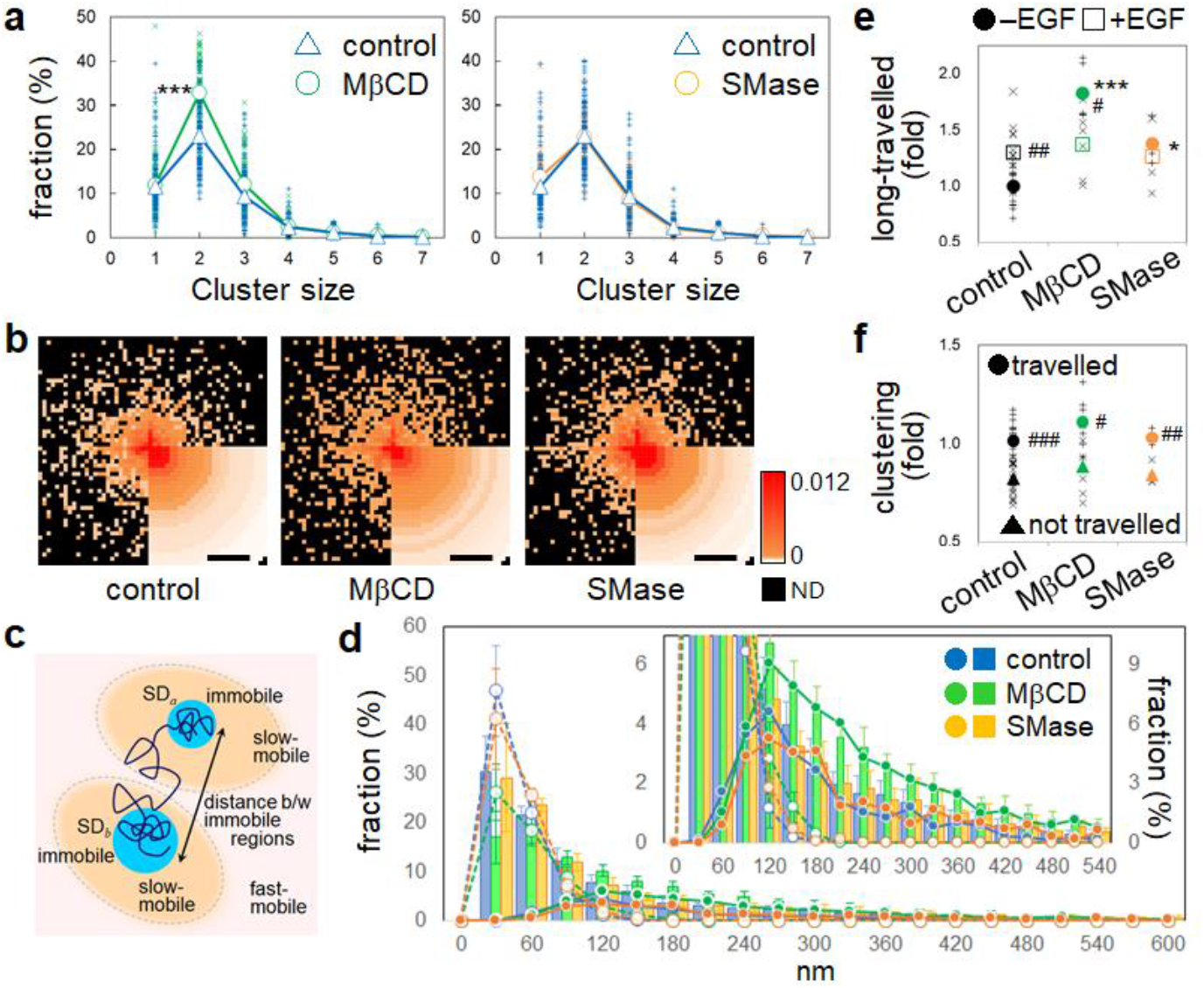
Lipid-depletion and EGFR pre-dimerization. **a**. Fractions of clusters belonging to the slow-mobile state. Averages (circles and triangles) and single-cell data (+, ×) are shown. **b**. Locations of pre-dimer formation of the mobile (slow and fast) EGFR molecules relative to the center of the first immobile region in each trajectory. The color code indicates the number of events (trajectory^-1^ s^-1^). Scale bars, 500 nm. **c**. A schematic illustration of an EGFR trajectory. Movements longer than the distance between two immobile regions are called “long-traveled”. **d**. Distribution of the traveling distances during 1.5 sec (circles; left scale) and normalized frequencies of pre-dimer formation (bars; right scale) during the slow- and fast-mobile states (percentage of total events). Dots indicate distributions for the long-traveled EGFR molecules, and hollow circles indicate other molecules. Inset: the distributions with small fractions are magnified. **e**. Relative fractions of long-traveled EGFR moved out of an immobile region during the observation period. **f.** Fractions of EGFR molecules with clustering (≥ dimer) in the mobile state after moving out of an immobile region (-EGF). In **e** and **f**, the fractions are normalized to control cells. *** and * p < 0.001 and < 0.05 (t-test), respectively, between control and lipid-depleted conditions. ###, ##, and # p < 0.001, < 0.01, and < 0.05 (t-test), respectively, between before and after EGF stimulation.

We calculated the influx and efflux of monomers and pre-dimers from the immobile and fast-mobile states to the slow-mobile state based on the HMM analysis (Tables S1 and S2). In the control condition, a significant net efflux of monomers (0.27 ± 0.05% s^-1^) was observed from the slow-mobile state, but there was no significant influx or efflux of the dimers. Under cholesterol-depletion, influx and efflux were balanced both for monomers and dimers in the slow-mobile state, thus producing no net flux. Although the decreased efflux of the monomer fraction from the slow-mobile state under cholesterol-depletion can induce pre-dimer formation, considering the 11.5% fraction of the slow-mobile state, this effect is small within the reaction time (subsecond) of the dimerization and decomposition (see below). However, this decreased efflux increased the fraction of the slow-mobile state. Because 49% and 63% of pre-dimer formations were observed during the slow-mobile state in the control and cholesterol-depleted conditions, respectively, the increase in the slow-mobile fraction increased the fraction of pre-dimers in the cholesterol-depletion condition. Transitions in the motional states correlated neither with the pre-dimer formation nor the decomposition.

We also measured the reaction rate constants of dimerization and dimer decomposition. The 1st-order dimerization rate constants in the slow-mobile state, were calculated from the frequency of dimerization events and found to be 7.3 ± 0.5 s^-1^ (123 cells) in the control condition and 7.4 ± 0.4 s^-1^ (69 cells) in the cholesterol-depleted condition. The difference was not statistically significant. The decomposition rate constants were 7.6 ± 0.05 s^-1^ under the control condition and 6.1 ± 0.06 s^-1^ in the cholesterol-depleted condition, which was a significant difference (Table S3). The total density of fluorescent particles on the cell surface (1.4 ± 0.7 μm^-2^) was not affected by the cholesterol depletion. When this particle density was applied to the region for slow-mobile motion, the fractions of monomers and dimers in the slow-mobile state were converted into particle densities of 0.32 ± 0.02 μm^-2^ and 0.67 ± 0.04 μm^-2^ in the control condition, and 0.28 ± 0.02 μm^-2^ and 0.74 ± 0.04 μm^-2^ in the cholesterol-depletion condition and resulted in dissociation constants (K_d_) of 0.33 ± 0.02 μm^-2^ and 0.23 ± 0.02 μm^-2^, respectively. Here, the decrease in the dissociation rate constant, i.e., the increase in the stability of pre-dimers, mainly contributed to the increased pre-dimer formation under the cholesterol depletion. Furthermore, the destabilization of the EGFR clusters suggested above may have another cause to increase the pre-dimer fraction.

Next, we checked locations of the pre-dimer formation relative to the center of the first immobile region along the single-molecule trajectories. The frequency of dimerization events per area in the slow- and fast-mobile states was mapped in two dimensions and averaged over the circumference (Fig. 3b). Reflecting the release from confinement (Fig. 2b), the pre-dimerization locations spread further away when cholesterol was depleted (“MβCD” in Fig. 3b). Most of the molecules traveled to the next immobile region (long-traveled; Fig. 3c) and formed pre-dimers regardless of the lipid condition (Fig. 3d). In the absence of EGF, the long-traveled fraction was largest upon cholesterol-depletion (Fig. 3e). Dimerization and higher-order clustering also occurred more frequently in the long-traveled fraction (Fig. 3f). These results suggest that cholesterol-depletion spreads the pre-dimers and pre-clusters (Fig. S7a) of EGFR over a large region of the plasma membrane.

On the other hand, sphingomyelin-depletion caused no obvious change regarding the EGFR clustering, such as the fraction distribution (Fig. 3a and S6), reaction rate constants (Tables S1 and S2), location of the dimerization (Fig 3b, d, and f), or fraction of long-traveled molecules (Fig. 3d).

### Cholesterol- or sphingomyelin-depletion inhibited EGF-induced clustering of EGFR

We quantitatively described the extent of EGF-induced clustering as the ratio of cluster fractions before and after EGF stimulation (Figs. 4a and S8a) and noticed obvious effects of the lipid depletion in the immobile state. In the control condition, the ratio for the same sized clusters larger than dimers was increased up to 7.0-fold, indicating that EGF stimulation reinforced higher-order clustering. When either cholesterol or sphingomyelin was depleted, this increase was strongly inhibited (only up to 2.5-fold), indicating that lipids facilitate the EGF-induced cluster formation. At the same time, the EGF-induced expansion of the locations of clustering (≥ trimer) was observed under the control condition but not under lipid-depletion (Fig. 4b). However, the area of the EGF-induced dimerization was similar under all lipid conditions (Fig. S7b). The EGF-induced dimerization during long traveling occurred at the same frequency between all lipid conditions (Fig. 4c), whereas higher-order clustering was facilitated by EGF only in the control condition (Fig. 4d) in parallel with the increase in long-traveled molecules (Fig. 3e).

**Fig. 4.**
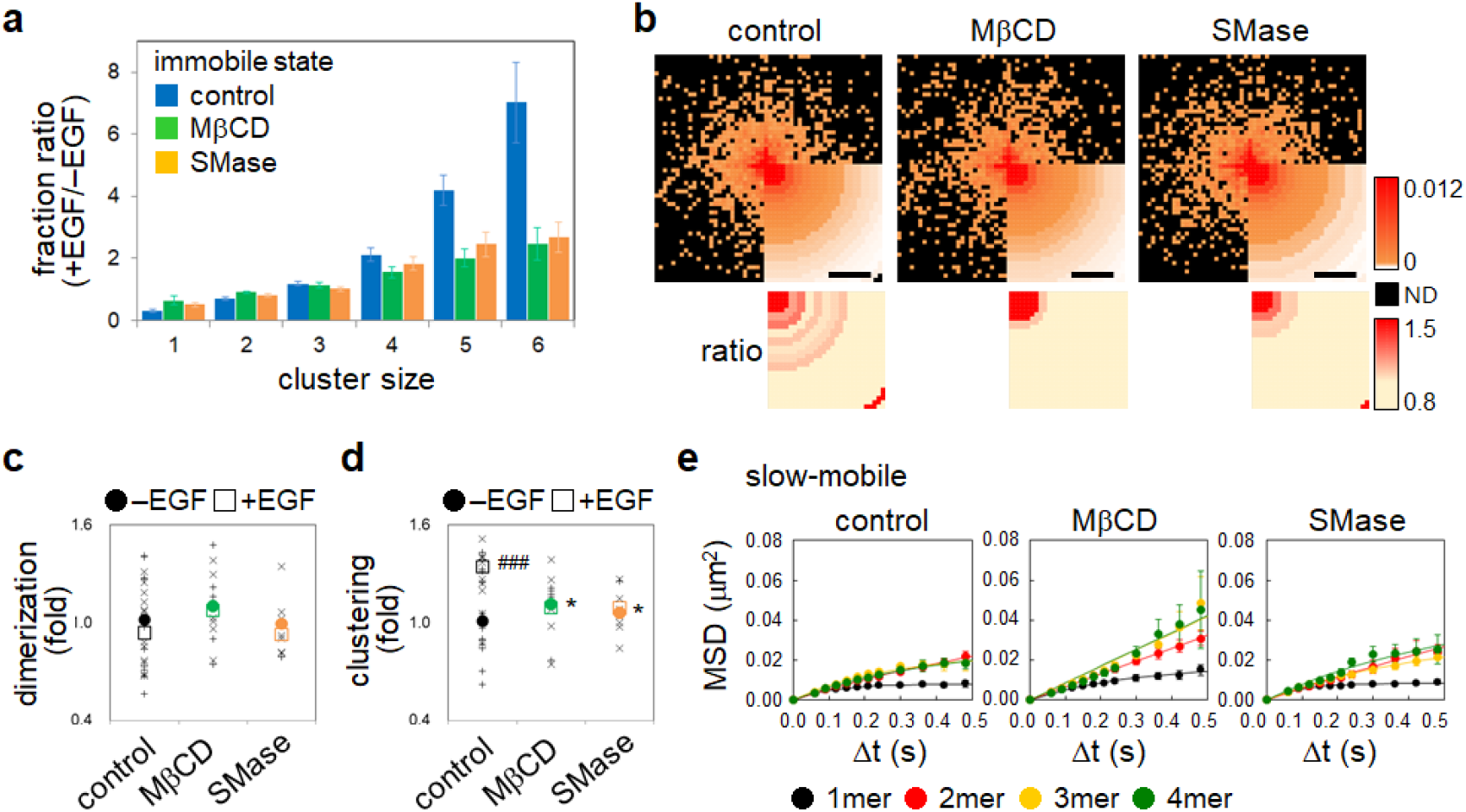
Lipid-depletion and EGF induced changes in mobility and clustering. **a**. The ratio of immobile clusters before and after EGF stimulation. Means and SE from 153 and 25 cells for control, 69 and 21 cells for cholesterol-depletion, and 50 and 27 cells for sphingomyelin-depletion are shown before and after the EGF stimulation, respectively. **b**. Top, locations of EGF-induced higher-order clustering during the mobile state (trajectory^-1^ s^-1^) relative to the center of the first immobile region. Bottom, ratios of clustering (after:before the EGF addition). Scale bars, 500 nm. **c** and **d**. Dimerization (**c**) and higher-order clustering (≥ trimer; **d**) events as the relative fraction among the total long-traveled molecules. The fractions are normalized to control cells before the EGF stimulation. * p < 0.05 (t-test) between control and lipid-depleted conditions; ### p < 0.001 (t-test) between before and after EGF stimulation. **e**. MSD-Δt plots of the slow-mobile state. Error bars: SE. All single-cell data points are shown in Fig. S4.

Changes in the cluster size distributions (Fig. 4a and S8) indicate that EGFR molecules were immobilized at the same time as higher-order clustering in the control condition. We previously reported that EGF stimulation of cells transiently releases the confinement of the slow-mobile diffusion at the very early stage (~30 sec), then shrinks the area of the confined diffusion of EGFR at the early stage (1~2 min). EGFR clusters were formed during this biphasic mobility change (Hiroshima et al., 2018), which depends on the EGF concentration (Yasui et al., 2018). Our observations in the present study confirmed that EGF induces clustering of long-traveled particles and shrinks the area for the slow-mobile state under the control and sphingomyelin-depleted conditions (Fig. 4e and S4) and reduced the confinement lengths to ~160 (monomer) and ~340 nm (≥ dimer) and ~160 (monomer) and ~530 nm (≥ dimer), respectively (Table S1). Even for molecules in the slow-mobile state moving without confinement under cholesterol-depletion, the diffusion coefficient and MSD were significantly decreased by EGF. The EGF-induced reduction in EGFR mobility was independent of higher-order clustering and not regulated by cholesterol or sphingomyelin.

### Cholesterol- or sphingomyelin-depletion inhibited signaling to adaptor proteins

The adaptor proteins GRB2 and SHC receive signals from activated EGFR. Their SRC homology 2 (SH2) domains interact with the phosphorylated tyrosine residues of EGFR (Lowenstein et al., 1992), causing them to translocate to the plasma membrane (Hiroshima et al., 2018; Yoshizawa et al., 2021) and result in the tyrosine phosphorylation of SHC. We conducted single-molecule imaging of GRB2-HaloTag labeled with tetramethylrhodamine (TMR) and a Western blotting analysis of SHC (p52SHC) phosphorylation in EGFR-GFP expressing cells. The number of GRB2 molecules on the plasma membrane (Fig. 5a and S9a) increased after EGF stimulation 2.5-fold in the control condition. Under cholesterol- and sphingomyelin-depletion, the increase of translocated GRB2 molecules was 1.0- and 1.5-fold, respectively. After cholesterol- and sphingomyelin-depletion, the EGF-induced phosphorylation of SHC Tyr 317 residue (Fig. 5b and S9b) was decreased to 0.73 ± 0.07- and 0.78 ± 0.08-fold, respectively, of the control condition (Fig. 5c). These reductions of EGFR/GRB2 and EGFR/SHC interactions by cholesterol-depletion were in contrast to the upregulated EGFR phosphorylation (Fig. 1).

**Fig. 5.**
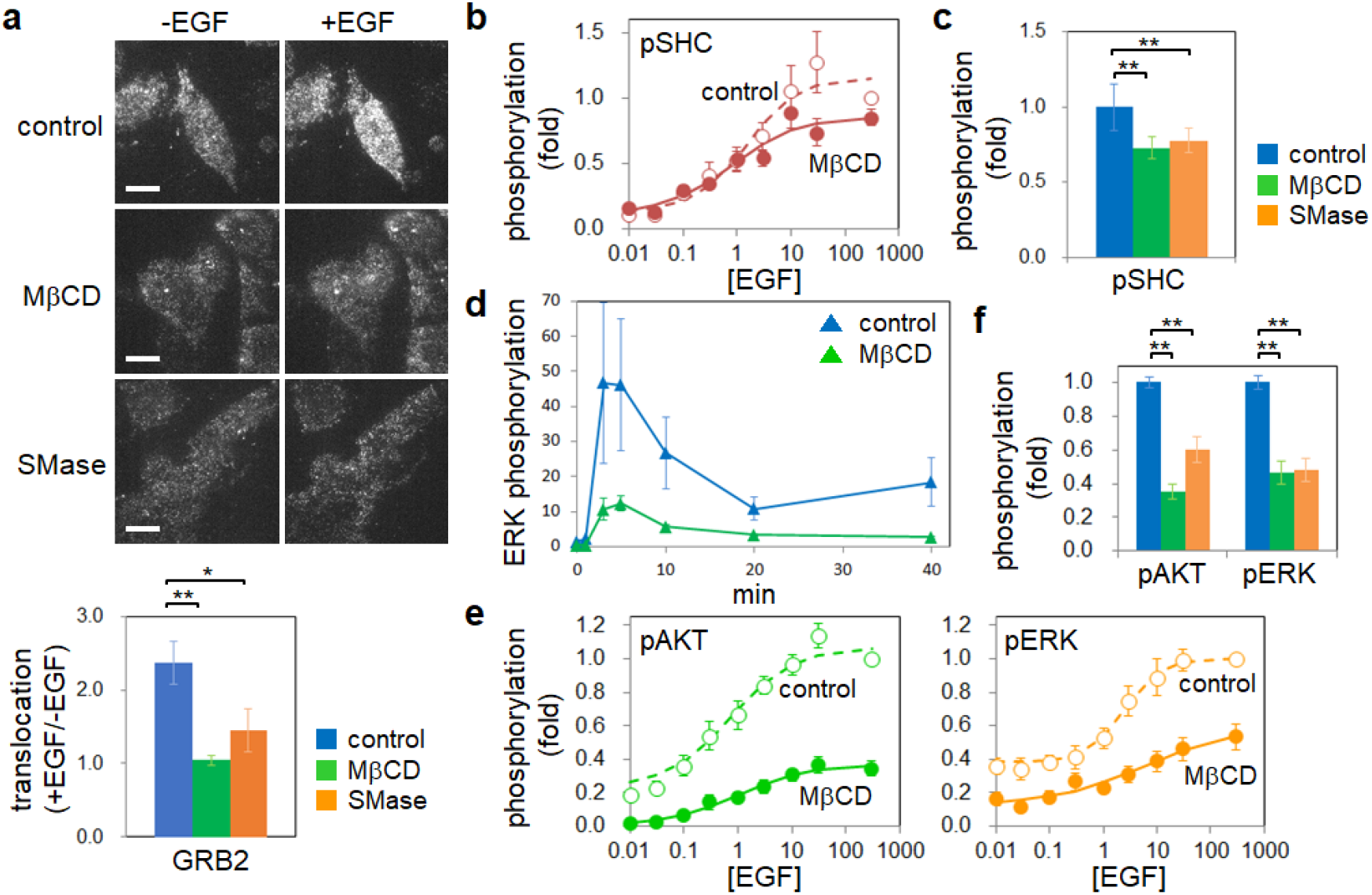
Lipid-depletion and downstream signaling. **a**. Translocation of GRB2. Top, single-molecule images of GRB2-HaloTag::TMR on the plasma membrane. Scale bars: 10 μm. Bottom, the ratio of single-molecule brightness on the plasma membrane in the same cells before and after the EGF stimulation (average of 6-8 trials). **b**. Dose-response curves for the EGF-induced phosphorylation of SHC Y317 (average of 6 trials). **c**. Phosphorylation levels of SHC at 30 nM EGF. **d**. Time-course of the EGF-induced ERK phosphorylation (average of 3 trials). Fold-changes of the phosphorylation level relative to 0 min. **e**. Dose-response curves for EGF-induced phosphorylation of downstream proteins (average of 7 and 13 trials for AKT and ERK, respectively). **f**. EGF-induced phosphorylation levels of AKT and ERK. * and ** p < 0.05 and p < 0.01, respectively. Error bars: SE. All data points are shown in Fig. S2b, S8, S9b, and S9c. In **a**, **b**, and **e**, the phosphorylation was measured 2 min after 30 nM EGF stimulation.

### Cholesterol- or sphingomyelin-depletion inhibited ERK and AKT phosphorylation

The phosphorylation of the downstream protein ERK in the EGFR signaling was measured by time-course Western blotting and maximized 5 minutes after the EGF stimulation, which is later than the time of EGFR phosphorylation (Figs. 5d and S10a). EGF-induced ERK phosphorylation was observed in all conditions, but its level was lower with cholesterol-depletion, which is expected when considering the reduced GRB2 translocation to the plasma membrane (Fig. 5a) and SHC phosphorylation (Fig. 5b and c). Unlike the adaptor proteins, the reduction of EGF-induced phosphorylation was remarkable in ERK under cholesterol-depletion. The dependency on EGF concentration was quantified for the phosphorylation of ERK and AKT, another downstream protein, 2 minutes after the stimulation (Figs. 5e and S10b). The EC_50_ values for the control and cholesterol-depleted conditions, which were obtained by fitting the dose-response curve (see Methods), were almost the same (within ±1 nM). The effects of the lipid depletions on the phosphorylation of AKT and ERK were evaluated 2 minutes after 30 nM EGF stimulation, which is almost the saturation condition (Figs. 5f and S10b). Although cholesterol-depletion increased the level of EGFR phosphorylation (Fig. 1e), the levels of AKT and ERK phosphorylation were significantly decreased (Fig. 5f). Sphingomyelin-depletion also lowered the levels of ERK and AKT phosphorylation, but differently, likely reflecting the specificities of the signaling pathways.

## Discussion

Cholesterol and sphingomyelin are major components of the plasma membrane subdomain where EGFR has been known to accumulate. We investigated how the depletion of either component affects EGFR behavior and signaling. We confirmed that the depletion of one lipid on the plasma membrane does not affect the other (Fig. S1) (Abe et al., 2012). Therefore, the observed phenomena in the present study, including downstream signaling of EGFR, were assumed to reflect the effects of each lipid.

EGFR phosphorylation in the early stage of EGF stimulation was upregulated under cholesterol-depletion (Figs. 1d and e). We considered this upregulation to depend on the increased amount of EGFR pre-dimer, which hardly undergoes auto-phosphorylation but is primed for a rapid response upon EGF stimulation (Teramura et al., 2006). EGFR has three motional modes in its lateral diffusion coefficient. After cholesterol-depletion, the amount of pre-dimer increased approximately 1.4-fold primarily in the slow-mobile state. The diffusion mode of the slow-mobile state was altered from confined to simple diffusion (Fig. 2b) without a significant change in the diffusion coefficient (Fig. S3a). This observation indicates that cholesterol-depletion enabled molecules to go freely through some barrier and move long distance (~1.8 fold longer than the control during the observation time). This barrier might correspond to a physical factor that maintains spatial phase separation in the membrane to impede EGFR from moving over the subdomain border composed of cholesterol or some component interacting with cholesterol around EGFR (e.g. shell model) (Anderson, 2002). Our results suggest that EGFR molecules in the slow-mobile state prefer to exist in the subdomains. Since EGFR pre-dimers were mainly present in the slow-mobile state, the disappearance of the barrier allowed them to spread over the cell surface.

The effect of cholesterol depletion on the affinity between EGFR protomers in the pre-dimer was considered from the rate constants of dimerization and decomposition. In the slow-mobile state, the rate constant of decomposition was significantly decreased, but we did not detect a significant change in the dimerization rate constant. As a result, the affinity was increased by the cholesterol depletion. In addition, the fraction of the slow-mobile state was increased due to increased and decreased of the transition probabilities (rate constants) from the fast-to slow-mobile states and the slow-to fast-mobile states, respectively (Table S1). These two effects induced the increase in the number of slow-mobile pre-dimers under the cholesterol-depleted condition and possibly resulted in the upregulation of EGFR phosphorylation. The disappearance of the diffusion barrier for the slow-mobile state of EGFR molecules may be related to the increase of the slow-mobile fraction. It also seems likely that the stimulative effect of cholesterol-depletion on the EGFR phosphorylation (Fig. 1) was caused by the stabilization of a pre-dimer structure for the kinase activation. Recently, we found that a transmembrane (TM)-juxtamembrane (JM) peptide of EGFR forms distinct structures of dimers in nanodiscs with or without cholesterol (Maeda et al., 2021). Cholesterol suppressed the formation of the JM dimer, which can be attributed to the structure suggested for EGFR kinase activation (Arkhipov et al., 2013). On the other hand, cholesterol stabilized dimers and trimers of EGFR peptides with lower JM interactions in the nanodiscs (Maeda et al. 2021).

Sphingomyelin-depletion, which did not affect cholesterol, also caused significant effects on EGFR in the slow-mobile state. The confinement length for the slow-mobile state was increased, though the confinement did not disappear (Fig. 2b). The fraction of the slow-mobile state was increased (Fig. S3b), reflecting the rise in the transition probability from the fast-to slow-mobile states (Table S1). The fractions of the monomer and other clusters in the slow-mobile state were unchanged. Sphingomyelin-depletion is likely to disrupt PIP_2_ domains, which locate at the cytoplasmic side of the sphingomyelin domains on the extracellular side of the plasma membrane (Abe et al., 2012). PIP_2_ facilitates dimer formation of the JM region of EGFR (Arkhipov et al., 2013; Maeda et al., 2018), and disruption of the PIP_2_ domain can cause EGFR monomerization. This disruption may be the reason for the unchanged pre-dimer fraction despite the increase in the slow-mobile state.

EGFR clusters larger than dimers were also formed before the EGF stimulation (pre-clusters). Different from pre-dimers, cholesterol-depletion did not increase the pre-cluster fraction (Fig. 3a), although the confinement disappeared (Fig. 2b) to enlarge the regions of the slow-mobile motions for the pre-clusters (Fig. 2c). On the contrary, the EGF-induced formation of higher-order clusters, which was observed in the control condition, was suppressed under cholesterol-depletion (Fig. 4a and 4b) in a cholesterol dose-dependent manner (Fig. S8b). Sphingomyelin-depletion also suppressed the EGF-induced clustering of EGFR. These lipid dependencies suggest that the clustering of EGFR is caused by a mechanism different from that for EGF-induced dimerization. Cholesterol and sphingomyelin may pack and enclose the EGFR molecules in small membrane subdomains or directly bind up the molecules. The oligomerization of TM peptides of EGFR has been observed in liposomes containing cholesterol (Jones et al., 1998). Following our previous report that the EGF-induced EGFR clusters in the immobile state are the primary interaction sites with the adaptor protein GRB2 (Hiroshima et al., 2018), the deficient clustering by the lipid depletion correlated with the reduction in the membrane translocation and in the phosphorylation of adaptor proteins (Fig. 5a and b). Indeed, the downstream proteins ERK and AKT showed reduced phosphorylation (Fig. 5c, 5d, and 5e), suggesting that cholesterol and sphingomyelin substantially contribute to the cellular signaling through the EGFR-immobile cluster formation.

Based on our observations (Fig. S11), we provide a scheme for EGFR-mediated cell signaling (Fig. 6): First, the immobile and slow-mobile states of EGFR are confined within a cholesterol- and sphingomyelin-enriched membrane subdomain (Fig. 2b). A significant fraction of EGFR molecules forms pre-dimers while moving within and between the slow-mobile state; however, only cholesterol and not sphingomyelin prevents the slow-mobile EGFR from freely passing over the border and interfering with the pre-dimer formation (Fig. 3a). Then, EGF association quickly converts EGFR from a pre-dimer to kinase active dimer. Moreover, EGF facilitates the formation of clusters larger than dimers with an enlarged area of the clustering (Fig. 4b and squares in 4d). This process involves the transient expansion of EGFR-cluster (not monomer) mobility at the very early stage of the EGF stimulation (Hiroshima et al., 2018). The clustering of EGFR requires cholesterol and sphingomyelin (Fig. 4a). Tyrosine phosphorylation of EGFR, a prerequisite for the clustering, leads to immobilization (Yasui et al., 2018), though the immobilization does not require the clustering. Finally, the immobile clusters increase and principally transduce information to the downstream cell signaling.

**Fig. 6.**
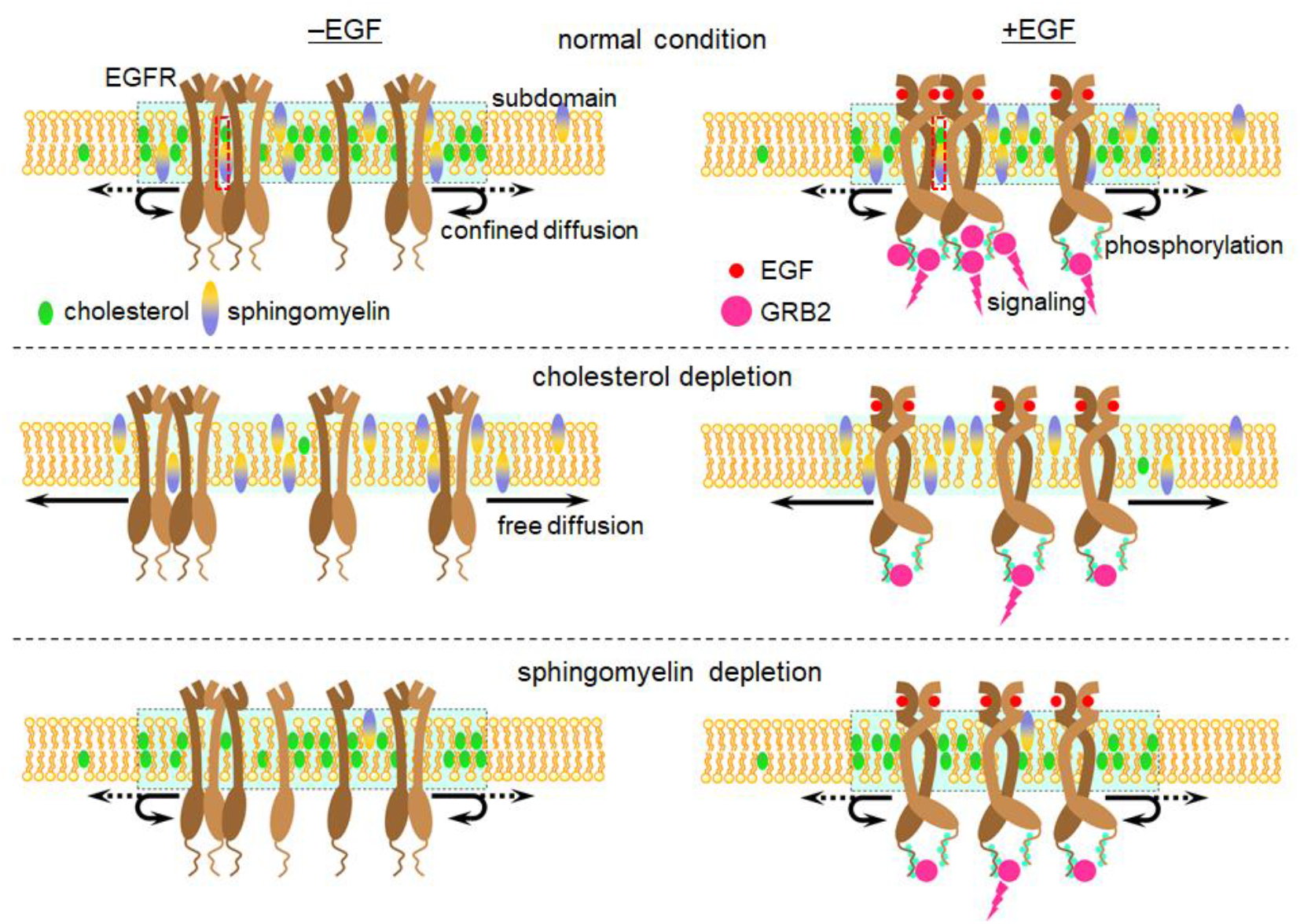
Lipid-depletion and EGFR signaling. Proposed EGFR dynamics in the slow-mobile state under control and lipid-depleted conditions. Dashed blue rectangles indicate the membrane subdomains that confine the EGFR mobility. Molecules clustered and immobilized during the slow-mobile state relay EGF-induced signaling depending on the lipid components.

Previous studies have reported both positive and negative effects of membrane cholesterol and lipid rafts in EGFR phosphorylation and downstream signaling (Chen and Resh, 2002; Fang et al., 2006; Liu et al., 2007; Zhuang et al., 2002). Here, we observed the dimerization and clustering of EGFR at single-molecule resolution and found a dichotomic effect of cholesterol, in which cholesterol suppresses the pre-dimerization of EGFR, leading to a reduction of EGF-induced phosphorylation, but assists with the EGF-induced higher-order clustering of phosphorylated EGFR to construct reaction sites for downstream signaling. This latter effect is common with sphingomyelin. The molecular mobility, dimerization/clustering, phosphorylation, and interaction with downstream molecules are intricately coupled in the process of EGFR signaling. Changes in the receptor behavior and membrane lipid environment can therefore cause variable results in the signal transduction, potentially causing the EGFR related diseases such as cell carcinomas, dyslipidemia, and so forth.

## Acknowledgment

We thank A. Yoshimura for the cDNA, H. Sato and A. Kanayama for experimental support, and P. Karagiannis for reading the manuscript. This study is supported by MEXT Japan with Grants-in-Aid for Scientific Research(B) (18H01839) and Grant-in-Aid for Scientific Research on Innovative Areas (18H05414). Y.S. was supported by MEXT Japan with Grants-in-Aid for Scientific Research (19H05647) and JST with CREST (JPMJCR1912).

## Author Contributions

M.H. and Y.S. designed the research; M.H. performed the experiments and analyzed the data; M.A. purified the fluorescence probes for lipids; M.A. and A.M. directed the lipid depletion study; N.T. and F. H-M quantified cellular cholesterol; M.U. and T.K. directed the study; and M.H. and Y.S. wrote the paper.

## Materials and Methods

### Gene Construction

The EGFR-GFP plasmid was constructed using the cDNA of human *EGFR* (*pNeoSRαII*) provided by Akihiko Yoshimura (Keio University) and was inserted into the pEGFP-C1 vector (Clontech) with the same linker sequence suggested by Carter and Sorkin (Carter and Sorkin, 1998). The GFP sequence included the monomeric mutation of A206K in the enhanced GFP (EGFP) sequence. GRB2-HaloTag was constructed using the human GRB2-encoding fragment and inserted into the Halo7-C2 vector in which the monomeric EGFP sequence in the pEGFP-C2 vector (Clontech) was substituted to the Halo7 sequence from the FN19K HaloTag T7 SP6 Flexi Vector (Promega).

### Cell Culture and Transfection

Chinese hamster ovary K1 (CHO-K1) cells were provided by RIKEN BRC through the National Bio-Resource Project (MEXT, Japan). For single-molecule and Western blotting experiments, a CHO cell line expressing EGFR-GFP was established. HAM F12 medium supplemented with 10% fetal bovine serum (FBS) was used to maintain the cells at 37 °C under 5% CO_2_.

### Cholesterol and sphingomyelin depletion

To deplete cholesterol, the cells were incubated in 5 or 10 mM MβCD (Sigma C4555) in Hank’s balanced salt solution (HBSS) for 1 hour at 37 °C under 5% CO_2_. Free cholesterol in the cells was separated by thin-layer chromatography (TLC) and quantified using gas chromatography-flame ionization detector (GC-FID, Shimadzu GC-14AH) or -mass spectrometry (GC/MS, JEOL JMS-700V). Cholesterol extent was also observed under a fluorescence microscope (Nikon, Ti) with a 20X objective lens (Nikon VC 20X, NA0.75) in θ toxin-GFP labeled cells. Sphingomyelin was depleted by incubating the cells in HBSS containing 1:300 diluted sphingomyelinase (Sigma, S9396) for 1 hour at 37 °C under 5% CO_2_. The depletion was confirmed by fluorescence microscopy in lysenin-GFP labeled cells. The average fluorescence intensity (per pixel) was measured over the cell region, and the averaged background intensity acquired from regions with no cells was subtracted from the signal.

### Microscopy and Image Analysis for Single-molecule Imaging and Tracking

Cell starvation was carried out by changing HAM F12 medium to modified Eagle’s medium minus phenol red and FBS 1 day before single-molecule imaging. Objective-type total internal reflection illumination was applied to observe EGFR-GFP in the basal plasma membrane of the cells through a PlanApo 60× NA 1.49 objective (Nikon, Tokyo, Japan) equipped on an inverted microscope (TE2000; Nikon). Lasers with wavelengths of 488 nm (Sapphire 488; Coherent, Santa Clara, CA) and 561 nm (Sapphire 561; Coherent) were used for the excitation of GFP and TMR, respectively. The dichroic mirror and emission filter were Di02-R488 (Semrock) and FF01-525/45 (Semrock) for GFP, and Di02-R561 (Semrock) and BLP02-561R (Semrock) for TMR imaging. An electron-multiplying CCD (EMCCD) camera (C9100-23; Hamamatsu, Hamamatsu, Japan), which was controlled using HCImage software, acquired fluorescence images at a frame rate of 33 s^-1^. The imaging was done at 25°C. Image processing was carried out with moving averages over two frames and background subtraction using rolling ball filtering (radius: 25 pixels) of the ImageJ plugins. Single-molecule tracking was performed on the processed images with custom-made software. The obtained data including positions and intensities of all fluorescent spots were analyzed using the methods described below.

### State Estimation Using a Hidden Markov Model with the Variational Bayes (VB-HMM) Method

A time series of step displacements and fluorescence intensities of the EGFR-GFP spots in the single-molecule tracking data were analyzed by VB-HMM analysis. This analysis consisted of the following steps (details are given in Okamoto and Sako, 2012; Persson et al., 2013). First, the data were grouped into *N* number of states with the K-means clustering method. Second, the initial parameters were calculated for each group based on observation probability models describing a two-dimensional diffusion equation for the step displacement and a Gaussian function for the fluorescence intensity. Third, the posterior probability distribution, *q*(*Z*, *θ*), where *Z* is the molecular state sequence and *θ* = {*π*, *A*, *ϕ*} is the parameters of the initial values, transition matrix, and the observation probability, respectively, was factorized as *q*(*Z*) *q*(*θ*). Then, the distribution functions, *q*(*Z*) and *q*(*θ*), were optimized with the VB expectation-maximization (VB-EM) algorithm. The VB-E and VB-M steps were alternately applied to optimize *q*(*Z*) with the forward-backward algorithm (Bishop, 2006) and *q*(*θ*) by updating the parameters, which were used in the next VB-E step. Fourth, the lower bound of the evidence, *L_q_*, was calculated to evaluate its convergence (except for the first *L_q_* value) by judging whether the difference from the previous *L_q_* was less than 0.001%. Fifth, if *L_q_* was not convergent, the next iteration was performed by repeating the third and fourth steps. Finally, the state sequence was determined by choosing the state with the highest probability at every frame.

### MSD for Each Mobility and Clustering State

The MSD of a specific mobility and clustering state, which was attributed to steps along the receptor trajectory, was calculated as 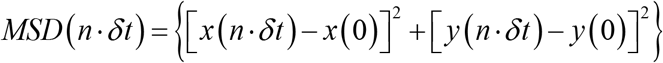, where *n* represents the frame number, *x* and *y* the particle positions, *δt* the time interval between frames (30 ms), and [] the average over the particles. By comparing the goodness of fit to equations for confined or simple diffusion using AIC (Eq. 4), the suitable diffusion model was determined for an MSD plot. The MSD calculation and all statistical tests were performed using Microsoft Excel.

### Translocation assay of adaptor proteins

Cell lines expressing both EGFR-GFP and GRB2-HaloTag were incubated in 96-well plates and starved 1 day before the experiment. The HaloTag-fused adaptor protein was labeled with 1-4 nM (depending on the GRB2 expression level) TMR and observed with 561-nm laser light for the excitation. For large-scale single-molecule analysis with high efficiency, well plate-based measurements were performed with the automated system that we developed (Yasui et al., 2018). Each of the automatically determined 5 fields of view, including 1-3 cells per field, was observed for 200 frames (6 sec) both before and 2 minutes after the EGF stimulation. The acquired images were analyzed using built-in software for tracking fluorescent spots. The spots observed in the 10th frame were used for the analysis to exclude fluorescence debris, which was bleached immediately after illumination. The number of translocated proteins on the plasma membrane was reflected in the total brightness of the fluorescent spots, in which more than one adaptor protein molecule might be included in a spot. The total brightness before and after the EGF stimulation were compared by their ratio.

### Western blot assay

The phosphorylation of proteins was quantified by western blotting using antibodies against pEGFR (#4407 for pY1173 and #3777 for pY1068; Cell Signaling Technology (CST)), pSHC (CST #2431 for pY317), pERK (CST #9106), and pAKT (CST #4060) to detect tyrosine or serine/threonine phosphorylation, and antibodies against EGFR (SC-03; Santacruz), SHC (CST #2432), ERK (CST #4696), and AKT (CST #9272) to detect protein expressions. Antibody binding was detected by luminescence using 1:2000 diluted HRP-linked anti-IgG antibodies (CST #7074 for rabbit and CST #7076 for mouse) as the secondary antibodies and ECL prime reagent (GE Healthcare). The luminescence intensities were measured using ImageJ software (NIH). Rectangular regions of interest were set in the signal (band) and background (far enough from the signal) regions. The difference in the average intensities of the two regions was defined as the band intensity. For the time-course analysis, the fold-change of the phosphorylation level at 0 min was calculated. The obtained band intensity at each time point in all conditions was normalized to that at 1 min of the control cells measured in the identical experiment. The intensity at 0 min was significantly weak, coupled with a relatively high level of noise, and not suitable as a normalization factor. Next, the normalized values at 0 min in all conditions were averaged and used as the denominator for the values at each time point. For the dose-response analysis, the obtained band intensity at each EGF concentration in all conditions was normalized to that at 300 nM EGF of the control cells measured in the identical experiment. The dose-response curve was fitted with the Hill equation as follows:

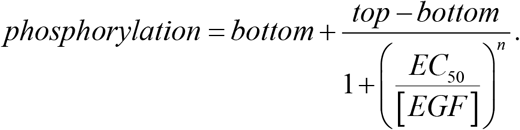

Here, *n*, *top*, and *bottom* are the fitted parameters indicating the Hill coefficient and upper and lower bounds, respectively. [*EGF*] is the concentration of EGF.

