## Supplementary Information for "Membrane cholesterol interferes with tyrosine phosphorylation but facilitates the clustering and signal transduction of EGFR"

### Fig. S1

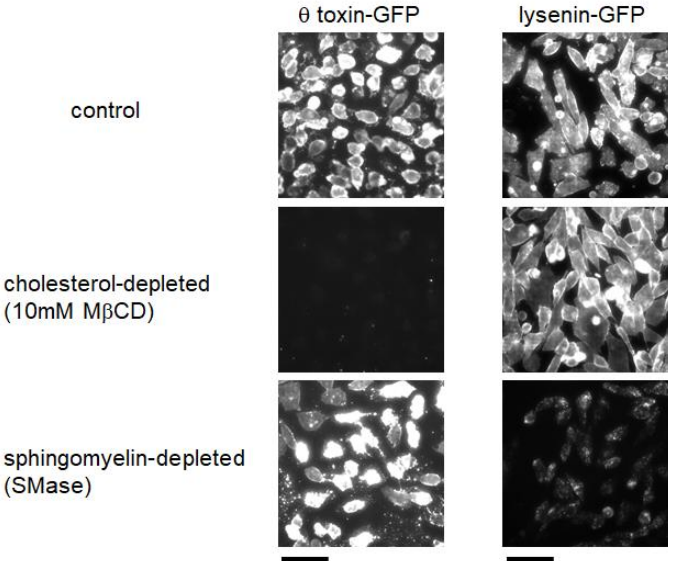


**Cholesterol- and sphingomyelin-depletion showed no mutual interference**. MβCD-, SMase-treated, and control CHO-K1 cells were labeled with probes of cholesterol (θ toxin-GFP) and sphingomyelin (lysenin-GFP). The cholesterol probe fluorescence was observed in the control and sphingomyelin-depleted cells, but not in the cholesterol-depleted cells. On the other hand, the sphingomyelin probe fluorescence was observed in the control and cholesterol-depleted cells, but not in the sphingomyelin-depleted cells. Scale bar, 50 μm.

**Fig. S2**

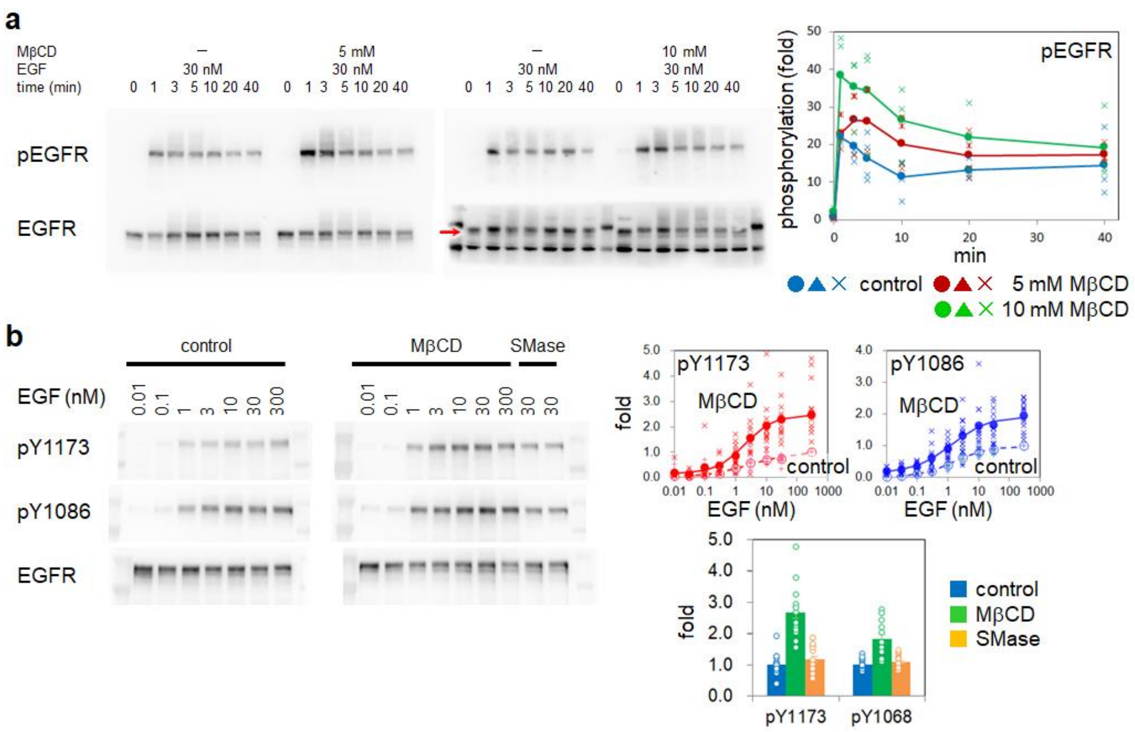


**Lipid-depletion and phosphorylation of signaling proteins**. **a**. Time-course western blotting with anti-phosphoEGFR (pY1173) antibody. EGFR-GFP was transfected in CHO-K1 (EGFR-null) cells. Blot images (left) and quantification results (right) are shown. The red arrow in **a** indicates the location of the pEGFR band. All obtained data points are shown. **b**. Left, western blots of pEGFR at the indicated EGF concentrations in the control and lipid-depleted conditions. Upper right, the dose-response curves for the cholesterol-depletion (10 mM MβCD) and control conditions. Lower right, comparison between the phosphorylation levels at 30 nM EGF under the indicated conditions. Circles show the result of independent measurements.

**Fig. S3**

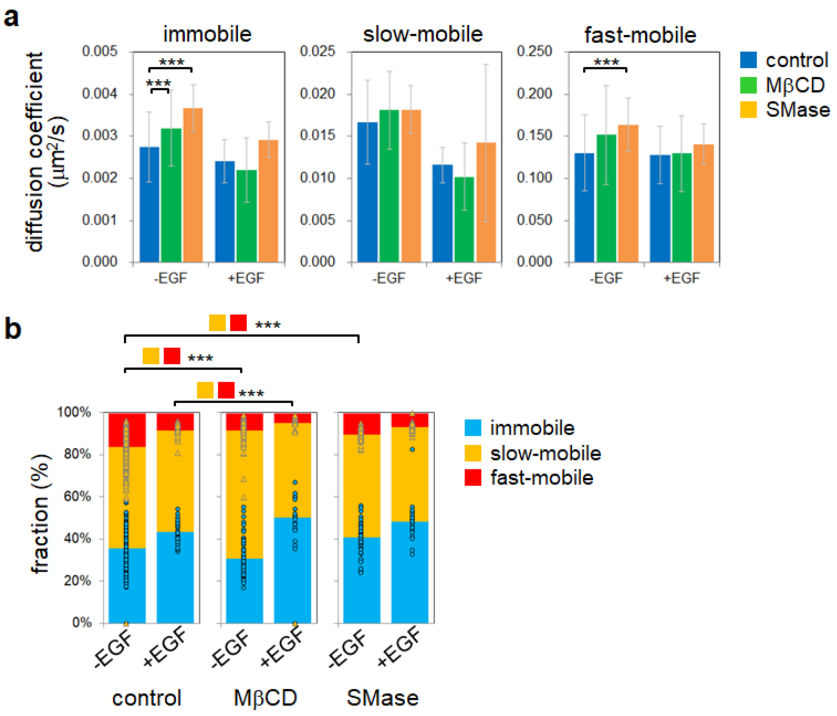


**Properties of EGFR motional states**. **a**. Diffusion coefficients and **b**. Fractions of each motional state under the indicated conditions. Cholesterol was depleted by 10 mM MβCD. *** $p < 0.001$ (t-test). 153 and 25 cells for the control, 69 and 21 cells for the cholesterol depletion, and 50 and 27 cells for the sphingomyelin depletion were measured before and after the EGF stimulation, respectively. Data in (**a**) are means and SD and in (**b**) are from single cells .

**Fig. S4**

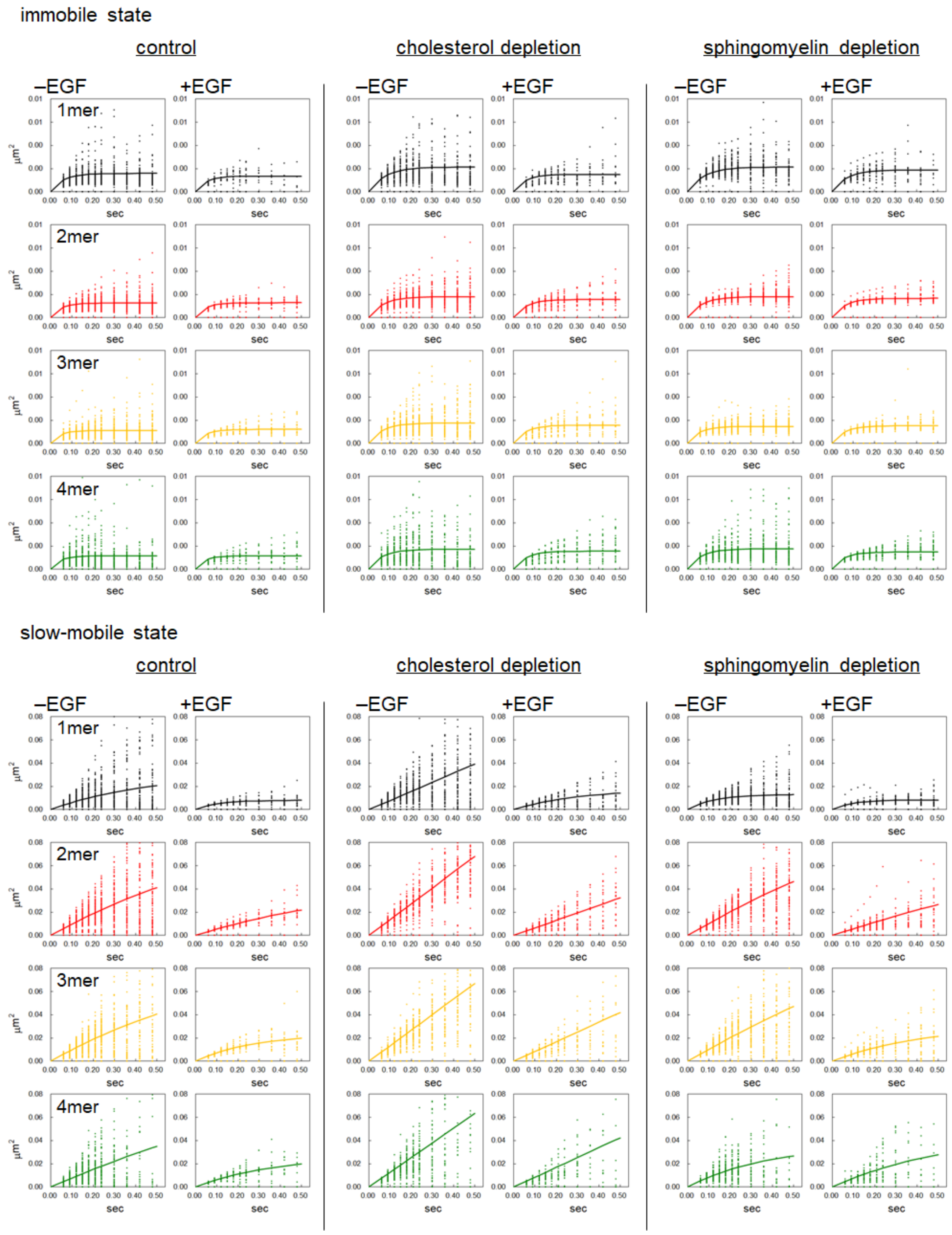
immobile state
control
cholesterol depletion
sphingomyelin depletion
–EGF
+EGF
1mer
2mer
3mer
4mer
µm²
sec
slow-mobile state
control
cholesterol depletion
sphingomyelin depletion
–EGF
+EGF
1mer
2mer
3mer
4mer
µm²
sec

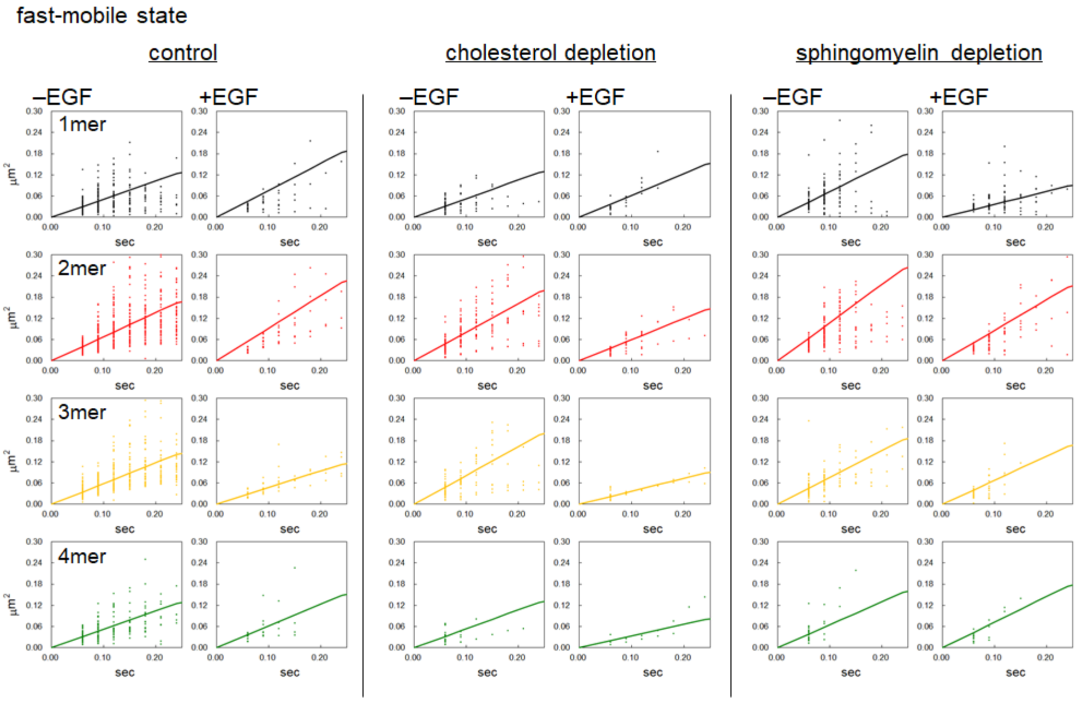


**EGFR mobility.** MSD values (points) obtained in single cells are shown as a function of lag times (x-axis). Cholesterol was depleted by 10 mM MβCD. Data from all observed cells are shown.

**Fig. S5**

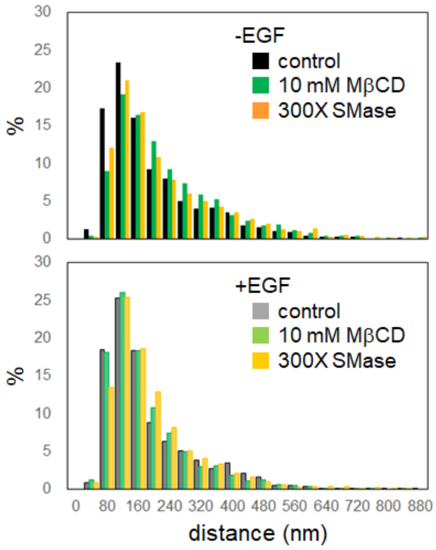


**Distance between immobile regions.** Distributions of the distances between the centers of the immobile regions observed in the EGFR trajectories are shown for control and lipid-depleted conditions with and without EGF stimulation.

**Fig. S6**

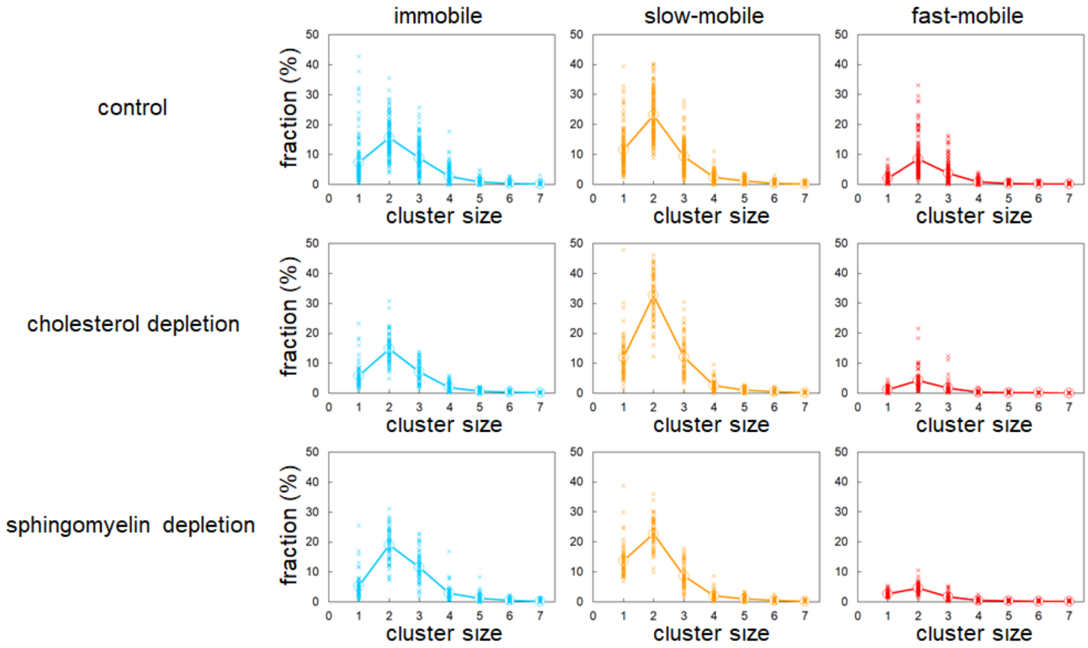


**Cluster-size distributions in the resting condition (without EGF stimulation).** Fractions of clusters in control, 10 mM MβCD-treated, and SMase-treated cells. All obtained data points are shown.

**Fig. S7**

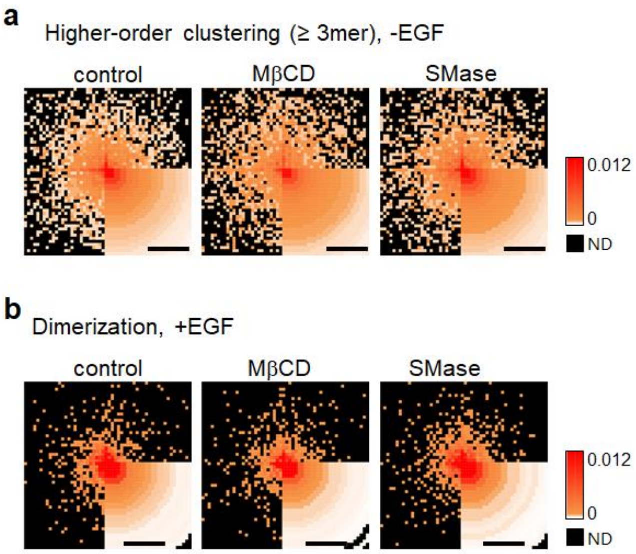


**Lipid-depletion and higher-order clustering**. **a**. Locations of EGFR higher-order pre-clustering (≥ trimer, -EGF condition) are plotted in a frequency (trajectory$^{-1}$ s$^{-1}$) heatmap relative to the center of the immobile region. **b**. Locations of EGF-induced EGFR dimerization relative to the center of the immobile region. Scale bars, 500 nm.

**Fig. S8**

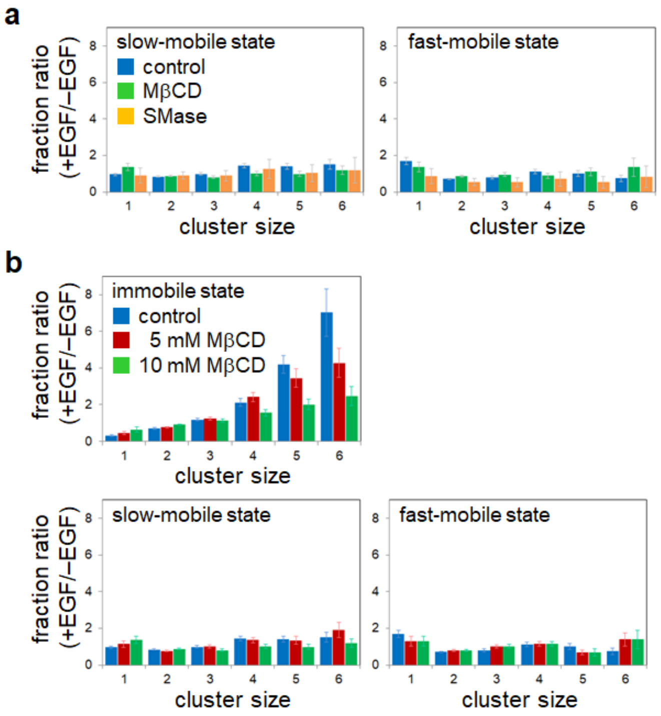


**Lipid-depletion and EGF-induced changes in the clustering**. The ratio of clusters before and after EGF stimulation. **a**. The ratios of the slow- and fast-mobile states in control, MβCD-, and SMase-treated cells. Those for the immobile state are shown in Fig. 4. **b**. Dependency on the MβCD concentration. Data show the mean and SD of 20-157 cells.

**Fig. S9**

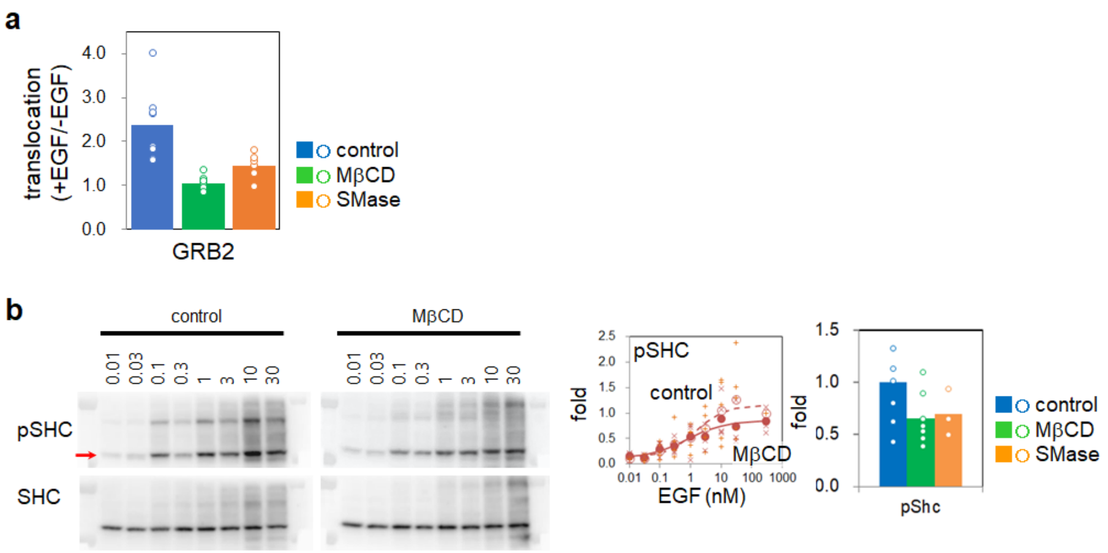


**Translocation to the plasma membrane and phosphorylation of adaptor proteins.** **a**. The brightness of GRB2-fluorescent spots summed over the plasma membrane. The ratios between total brightness before and after EGF stimulation in the same cells are shown. Each circle indicates the averaged ratio over 10-60 cells observed in 8 (control, cholesterol-depletion) or 6 (sphingomyelin-depletion) independent measurements. **b**. Left, western blots of phosphoSHC at the indicated EGF concentrations. Middle, the dose-response curves for the cholesterol-depletion (by 10 mM MβCD) and control conditions. Right, comparison between the phosphorylation levels at 30 nM EGF under the indicated conditions. Circles show the results of independent measurements.

**Fig. S10**

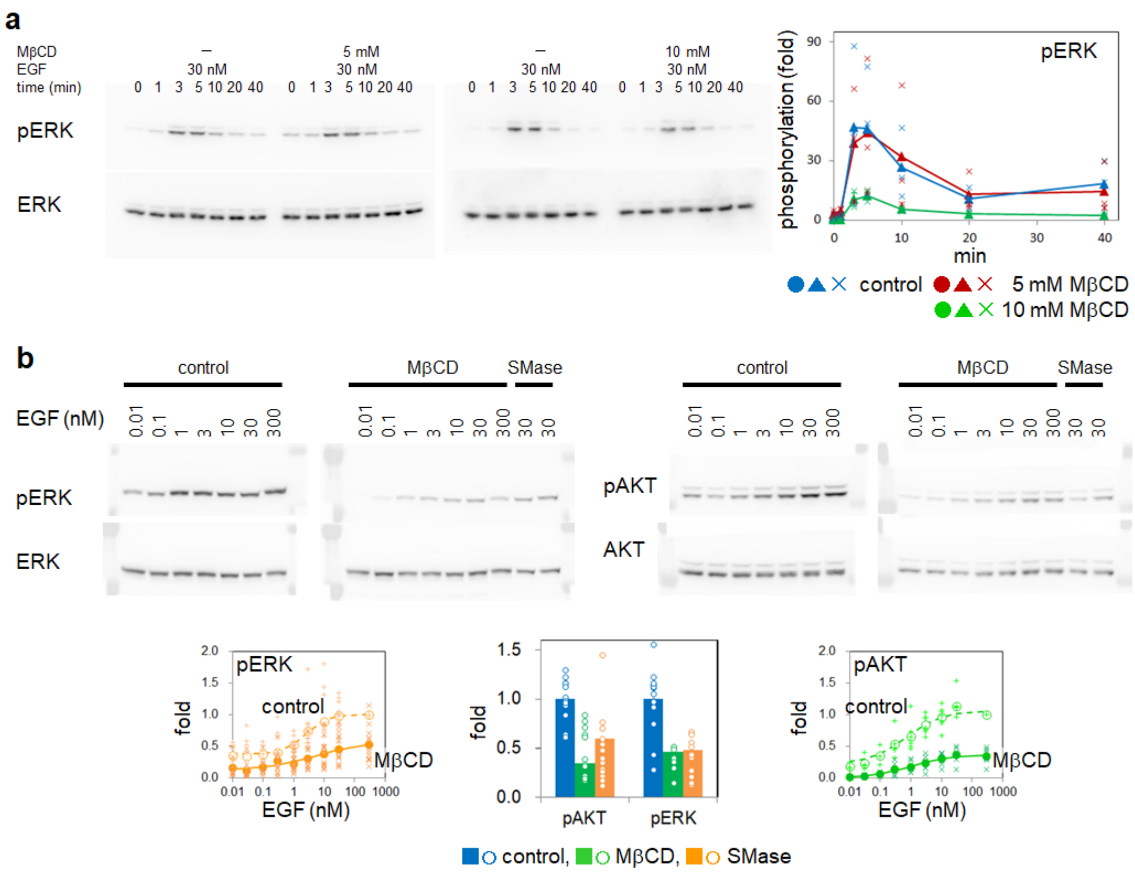


**Lipid-depletion and downstream signaling**. **a**, Time-course western blots using anti-phosphoERK antibody. Blots (left) and quantification results (right). All obtained data points are shown. **b**, EGF-dose-dependent phosphorylation of ERK (left) and AKT (right). Lower left and right, the dose-response curves of ERK (left) and AKT (right) for the control and cholesterol-depleted (by 10 mM MβCD) conditions, respectively. Lower middle, comparison between the phosphorylation levels at 30 nM EGF. All obtained data points are shown.

**Fig. S11**

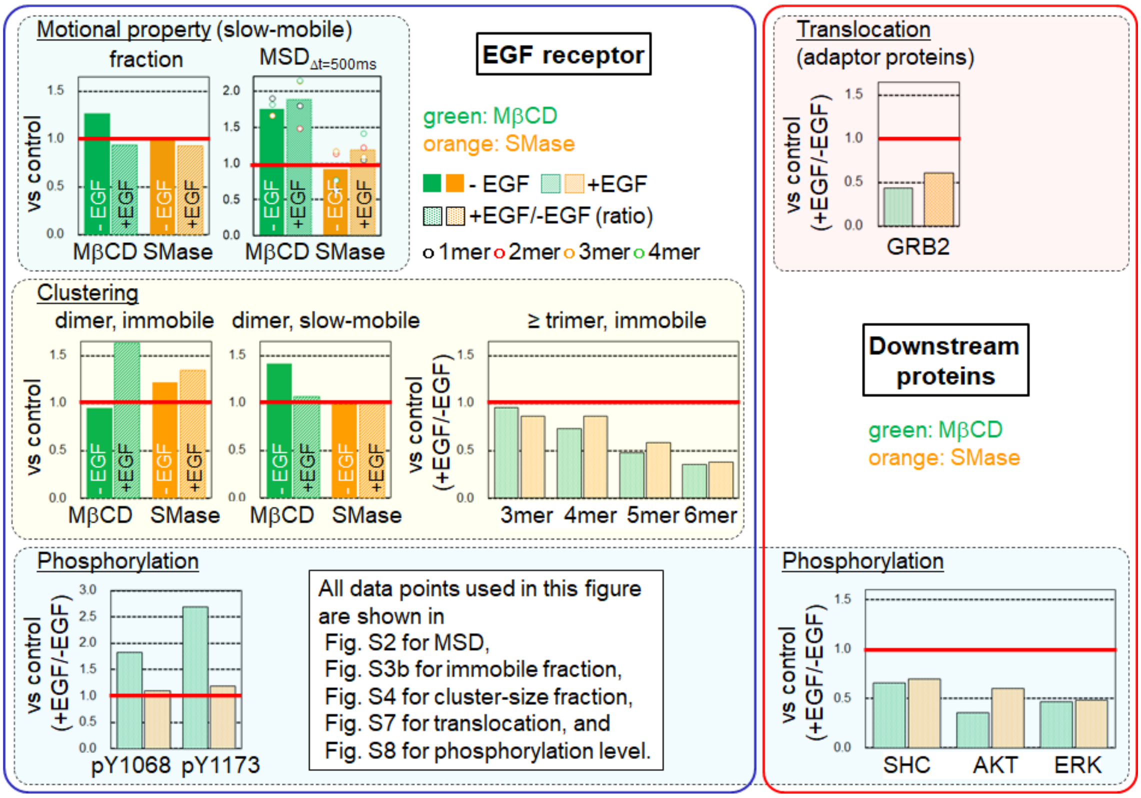


**EGF-induced changes in EGFR behavior and downstream signaling.** EGFR mobility (motional state fractions and MSD at a time lag of 500 ms), clustering (fractions), and signaling (phosphorylation levels) under lipid-depletion are compared with control. Data used for the calculation of the ratios are from the indicated figures. The EGFR mobility, dimerization, and phosphorylation in the lipid-depletion conditions exceeded the control level (red bars) except for higher-order clustering. On the other hand, the signaling to adaptor proteins that directly targets EGFR phosphorylation and the phosphorylation of downstream proteins was below the control level.

**Table S1 Parameters of EGFR motional states obtained by HMM analysis.**

| | | | -EGF | | | +EGF | | |
|---|---|---|---|---|---|---|---|---|
| control | | # of cells : | n= 152 | | | n= 25 | | |
| | Diffusion coefficient ($\mu m^2/s$) | immobile | 0.003 ± 0.001 | | | 0.002 ± 0.001 | | |
| | | slow-mobile | 0.017 ± 0.005 | | | 0.012 ± 0.002 | | |
| | | fast-mobile | 0.130 ± 0.046 | | | 0.128 ± 0.034 | | |
| | Diffusion mode | immobile | confined ( 60 nm ) | | | confined ( 60 nm ) | | |
| | | slow-mobile | confined ( 570 nm for ≥2mer ) | | | confined ( 340 nm for ≥2mer ) | | |
| | | fast-mobile | free | | | free | | |
| | Fraction (%) | immobile | 36 ± 9 | | | 44 ± 5 | | |
| | | slow-mobile | 48 ± 9 | | | 48 ± 4 | | |
| | | fast-mobile | 16 ± 8 | | | 8 ± 3 | | |
| | Transition array (i →j) | i \ j | immobile | slow-mobile | fast-mobile | immobile | slow-mobile | fast-mobile |
| | | immobile | 0.93 ± 0.02 | 0.07 ± 0.02 | 0.00 ± 0.00 | 0.92 ± 0.02 | 0.07 ± 0.02 | 0.00 ± 0.00 |
| | | slow-mobile | 0.05 ± 0.02 | 0.79 ± 0.04 | 0.15 ± 0.04 | 0.07 ± 0.01 | 0.84 ± 0.03 | 0.09 ± 0.02 |
| | | fast-mobile | 0.01 ± 0.01 | 0.41 ± 0.16 | 0.57 ± 0.16 | 0.01 ± 0.01 | 0.44 ± 0.10 | 0.54 ± 0.10 |
| MβCD 10 mM | | | n= 69 | | | n= 20 | | |
| | Diffusion coefficient ($\mu m^2/s$) | immobile | 0.003 ± 0.001 *** | | | 0.002 ± 0.001 | | |
| | | slow-mobile | 0.018 ± 0.005 | | | 0.010 ± 0.004 | | |
| | | fast-mobile | 0.152 ± 0.059 | | | 0.130 ± 0.045 | | |
| | Diffusion mode | immobile | confined ( 70 nm ) | | | confined ( 70 nm ) | | |
| | | slow-mobile | free | | | free & confined | | |
| | | fast-mobile | free | | | free | | |
| | Fraction (%) | immobile | 31 ± 8 *** | | | 50 ± 8 | | |
| | | slow-mobile | 61 ± 8 *** | | | 45 ± 7 | | |
| | | fast-mobile | 8 ± 6 *** | | | 5 ± 2 *** | | |
| | Transition array (i →j) | i \ j | immobile | slow-mobile | fast-mobile | immobile | slow-mobile | fast-mobile |
| | | immobile | 0.91 ± 0.03 *** | 0.08 ± 0.02 *** | 0.01 ± 0.00 *** | 0.92 ± 0.01 | 0.07 ± 0.01 | 0.00 ± 0.00 |
| | | slow-mobile | 0.04 ± 0.02 | 0.86 ± 0.03 *** | 0.10 ± 0.04 *** | 0.08 ± 0.02 | 0.86 ± 0.02 | 0.06 ± 0.03 *** |
| | | fast-mobile | 0.03 ± 0.03 *** | 0.56 ± 0.12 *** | 0.42 ± 0.13 *** | 0.01 ± 0.01 | 0.48 ± 0.11 | 0.50 ± 0.11 |
| SMase | | | n= 61 | | | n= 27 | | |
| | Diffusion coefficient ($\mu m^2/s$) | immobile | 0.004 ± 0.001 *** | | | 0.003 ± 0.000 *** | | |
| | | slow-mobile | 0.018 ± 0.003 | | | 0.014 ± 0.009 | | |
| | | fast-mobile | 0.164 ± 0.032 *** | | | 0.141 ± 0.024 | | |
| | Diffusion mode | immobile | confined ( 70 nm ) | | | confined ( 70 nm ) | | |
| | | slow-mobile | confined ( 720 nm for ≥2mer ) | | | confined ( 530 nm for ≥2mer ) | | |
| | | fast-mobile | free | | | free | | |
| | Fraction (%) | immobile | 41 ± 7 *** | | | 48 ± 9 | | |
| | | slow-mobile | 49 ± 6 | | | 45 ± 7 | | |
| | | fast-mobile | 10 ± 3 *** | | | 7 ± 2 | | |
| | Transition array (i →j) | i \ j | immobile | slow-mobile | fast-mobile | immobile | slow-mobile | fast-mobile |
| | | immobile | 0.92 ± 0.03 | 0.08 ± 0.03 | 0.01 ± 0.00 | 0.92 ± 0.01 | 0.08 ± 0.01 | 0.00 ± 0.00 |
| | | slow-mobile | 0.07 ± 0.02 | 0.78 ± 0.03 | 0.15 ± 0.03 | 0.09 ± 0.04 | 0.82 ± 0.03 | 0.09 ± 0.03 |
| | | fast-mobile | 0.02 ± 0.01 *** | 0.53 ± 0.06 *** | 0.45 ± 0.07 *** | 0.02 ± 0.03 | 0.49 ± 0.03 | 0.49 ± 0.04 |

*** $p < 0.001$ in comparison with control.

### Table S2 Parameters of EGFR clustering states obtained by HMM analysis.

| | | | -EGF | | | | | | +EGF | | | | | |
|---|---|---|---|---|---|---|---|---|---|---|---|---|---|---|
| | | | | | | | # of cells : n= 152 | | | | | | | n= 25 |
| control | | | 1mer | 2mer | 3mer | 4mer | 5mer | 6mer | 1mer | 2mer | 3mer | 4mer | 5mer | 6mer |
| | Clusters (%) | immobile | 7.3 ± 7.0 | 15.6 ± 5.9 | 8.9 ± 4.8 | 2.7 ± 2.7 | 0.8 ± 0.8 | 0.3 ± 0.5 | 2.5 ± 1.5 | 13.4 ± 5.5 | 12.6 ± 3.7 | 6.5 ± 2.2 | 3.9 ± 1.7 | 2.3 ± 1.5 |
| | | slow-mobile | 11.5 ± 6.5 | 23.2 ± 7.5 | 9.3 ± 5.0 | 2.5 ± 1.6 | 1.1 ± 0.9 | 0.4 ± 0.6 | 12.0 ± 3.5 | 20.4 ± 2.5 | 9.8 ± 3.2 | 3.8 ± 1.6 | 1.7 ± 0.9 | 0.7 ± 0.6 |
| | | fast-mobile | 2.0 ± 1.4 | 8.5 ± 5.9 | 3.7 ± 3.5 | 0.8 ± 0.7 | 0.4 ± 0.4 | 0.2 ± 0.3 | 2.1 ± 0.9 | 3.3 ± 1.7 | 1.6 ± 1.5 | 0.5 ± 0.4 | 0.2 ± 0.2 | 0.1 ± 0.1 |
| | | i \ j | 1mer | 2mer | 3mer | 4mer | 5mer | 6mer | 1mer | 2mer | 3mer | 4mer | 5mer | 6mer |
| | Transition array (i →j) | 1mer | 0.85 ± 0.05 | 0.11 ± 0.04 | 0.01 ± 0.01 | 0.00 ± 0.00 | 0.00 ± 0.00 | 0.00 ± 0.00 | 0.84 ± 0.04 | 0.14 ± 0.03 | 0.01 ± 0.01 | 0.00 ± 0.00 | 0.00 ± 0.00 | 0.00 ± 0.00 |
| | | 2mer | 0.06 ± 0.03 | 0.87 ± 0.03 | 0.05 ± 0.02 | 0.01 ± 0.01 | 0.00 ± 0.00 | 0.00 ± 0.00 | 0.06 ± 0.02 | 0.87 ± 0.03 | 0.06 ± 0.02 | 0.00 ± 0.00 | 0.00 ± 0.00 | 0.00 ± 0.00 |
| | | 3mer | 0.01 ± 0.01 | 0.10 ± 0.03 | 0.83 ± 0.04 | 0.02 ± 0.01 | 0.01 ± 0.01 | 0.00 ± 0.00 | 0.01 ± 0.00 | 0.09 ± 0.02 | 0.85 ± 0.03 | 0.04 ± 0.01 | 0.01 ± 0.00 | 0.00 ± 0.00 |
| | | 4mer | 0.02 ± 0.02 | 0.05 ± 0.04 | 0.05 ± 0.03 | 0.81 ± 0.06 | 0.01 ± 0.01 | 0.01 ± 0.01 | 0.01 ± 0.01 | 0.01 ± 0.01 | 0.07 ± 0.02 | 0.85 ± 0.04 | 0.03 ± 0.02 | 0.01 ± 0.01 |
| | | 5mer | 0.02 ± 0.02 | 0.04 ± 0.03 | 0.05 ± 0.03 | 0.01 ± 0.01 | 0.81 ± 0.06 | 0.01 ± 0.00 | 0.02 ± 0.03 | 0.02 ± 0.01 | 0.02 ± 0.01 | 0.05 ± 0.02 | 0.84 ± 0.05 | 0.02 ± 0.02 |
| | | 6mer | 0.02 ± 0.03 | 0.03 ± 0.03 | 0.03 ± 0.03 | 0.02 ± 0.02 | 0.01 ± 0.00 | 0.80 ± 0.08 | 0.01 ± 0.02 | 0.02 ± 0.02 | 0.01 ± 0.01 | 0.02 ± 0.01 | 0.03 ± 0.02 | 0.85 ± 0.05 |
| | | | | | | | | n= 69 | | | | | | n= 20 |
| MβCD 10 mM | | | 1mer | 2mer | 3mer | 4mer | 5mer | 6mer | 1mer | 2mer | 3mer | 4mer | 5mer | 6mer |
| | Clusters (%) | immobile | 5.7 ± 4.1 | 14.9 ± 4.9 | 7.2 ± 3.3 | 2.0 ± 1.4 | 0.7 ± 0.6 | 0.3 ± 0.4 | 5.6 ± 5.0 | 21.9 ± 5.8 | 13.4 ± 4.7 | 4.9 ± 1.7 | 2.4 ± 1.2 | 1.3 ± 0.9 |
| | | slow-mobile | 11.9 ± 7.1 | 32.7 ± 8.0 | 12.1 ± 6.5 | 2.6 ± 1.7 | 1.1 ± 0.7 | 0.5 ± 0.5 | 12.4 ± 6.9 | 21.8 ± 6.0 | 7.5 ± 3.0 | 2.0 ± 1.0 | 0.9 ± 0.6 | 0.4 ± 0.3 |
| | | fast-mobile | 1.1 ± 0.9 | 4.3 ± 3.7 | 1.6 ± 2.1 | 0.4 ± 0.3 | 0.3 ± 0.2 | 0.1 ± 0.1 | 1.0 ± 0.8 | 2.1 ± 1.5 | 0.8 ± 0.5 | 0.2 ± 0.1 | 0.2 ± 0.1 | 0.1 ± 0.1 |
| | | i \ j | 1mer | 2mer | 3mer | 4mer | 5mer | 6mer | 1mer | 2mer | 3mer | 4mer | 5mer | 6mer |
| | Transition array (i →j) | 1mer | 0.84 ± 0.04 | 0.13 ± 0.03 | 0.01 ± 0.01 | 0.00 ± 0.00 | 0.00 ± 0.00 | 0.00 ± 0.00 | 0.85 ± 0.05 | 0.13 ± 0.04 | 0.01 ± 0.01 | 0.00 ± 0.00 | 0.00 ± 0.00 | 0.00 ± 0.00 |
| | | 2mer | 0.05 ± 0.02 | 0.88 ± 0.03 | 0.05 ± 0.02 | 0.00 ± 0.00 | 0.00 ± 0.00 | 0.00 ± 0.00 | 0.05 ± 0.01 | 0.89 ± 0.03 | 0.05 ± 0.02 | 0.00 ± 0.00 | 0.00 ± 0.00 | 0.00 ± 0.00 |
| | | 3mer | 0.01 ± 0.01 | 0.11 ± 0.02 | 0.84 ± 0.03 | 0.02 ± 0.01 | 0.01 ± 0.01 | 0.00 ± 0.00 | 0.01 ± 0.00 | 0.09 ± 0.02 | 0.86 ± 0.03 | 0.03 ± 0.01 | 0.01 ± 0.00 | 0.00 ± 0.00 |
| | | 4mer | 0.02 ± 0.01 | 0.04 ± 0.04 | 0.06 ± 0.03 | 0.81 ± 0.05 | 0.01 ± 0.01 | 0.01 ± 0.01 | 0.01 ± 0.00 | 0.01 ± 0.01 | 0.06 ± 0.02 | 0.86 ± 0.03 | 0.03 ± 0.02 | 0.01 ± 0.00 |
| | | 5mer | 0.02 ± 0.03 | 0.04 ± 0.03 | 0.05 ± 0.03 | 0.02 ± 0.01 | 0.80 ± 0.07 | 0.01 ± 0.00 | 0.02 ± 0.05 | 0.02 ± 0.02 | 0.02 ± 0.01 | 0.04 ± 0.02 | 0.85 ± 0.06 | 0.01 ± 0.01 |
| | | 6mer | 0.01 ± 0.02 | 0.03 ± 0.03 | 0.02 ± 0.02 | 0.02 ± 0.02 | 0.01 ± 0.01 | 0.81 ± 0.07 | 0.02 ± 0.02 | 0.02 ± 0.02 | 0.02 ± 0.01 | 0.02 ± 0.01 | 0.02 ± 0.01 | 0.84 ± 0.06 |
| | | | | | | | | n= 61 | | | | | | n= 27 |
| SMase | | | 1mer | 2mer | 3mer | 4mer | 5mer | 6mer | 1mer | 2mer | 3mer | 4mer | 5mer | 6mer |
| | Clusters (%) | immobile | 5.3 ± 4.2 | 19.0 ± 5.6 | 11.7 ± 4.9 | 2.9 ± 2.5 | 1.2 ± 1.4 | 0.5 ± 0.5 | 3.2 ± 1.8 | 18.1 ± 5.2 | 14.3 ± 4.5 | 6.1 ± 1.8 | 3.3 ± 1.5 | 1.6 ± 1.2 |
| | | slow-mobile | 13.8 ± 5.5 | 22.8 ± 5.0 | 8.6 ± 4.0 | 2.0 ± 1.5 | 1.0 ± 0.9 | 0.5 ± 0.5 | 12.5 ± 2.7 | 20.6 ± 4.3 | 7.7 ± 2.4 | 2.5 ± 1.0 | 1.0 ± 0.5 | 0.5 ± 0.5 |
| | | fast-mobile | 2.7 ± 0.9 | 4.5 ± 1.6 | 1.6 ± 1.2 | 0.5 ± 0.4 | 0.3 ± 0.3 | 0.2 ± 0.2 | 2.3 ± 0.8 | 2.5 ± 1.0 | 0.9 ± 0.4 | 0.3 ± 0.2 | 0.1 ± 0.1 | 0.1 ± 0.1 |
| | | i \ j | 1mer | 2mer | 3mer | 4mer | 5mer | 6mer | 1mer | 2mer | 3mer | 4mer | 5mer | 6mer |
| | Transition array (i →j) | 1mer | 0.84 ± 0.03 | 0.12 ± 0.02 | 0.01 ± 0.01 | 0.01 ± 0.01 | 0.00 ± 0.00 | 0.00 ± 0.00 | 0.84 ± 0.03 | 0.13 ± 0.02 | 0.01 ± 0.01 | 0.01 ± 0.00 | 0.00 ± 0.00 | 0.00 ± 0.00 |
| | | 2mer | 0.06 ± 0.02 | 0.87 ± 0.02 | 0.05 ± 0.02 | 0.01 ± 0.01 | 0.00 ± 0.00 | 0.00 ± 0.00 | 0.06 ± 0.01 | 0.87 ± 0.02 | 0.06 ± 0.02 | 0.00 ± 0.00 | 0.00 ± 0.00 | 0.00 ± 0.00 |
| | | 3mer | 0.01 ± 0.01 | 0.09 ± 0.02 | 0.86 ± 0.03 | 0.02 ± 0.01 | 0.01 ± 0.01 | 0.00 ± 0.00 | 0.01 ± 0.00 | 0.10 ± 0.02 | 0.85 ± 0.03 | 0.03 ± 0.01 | 0.00 ± 0.00 | 0.00 ± 0.00 |
| | | 4mer | 0.02 ± 0.03 | 0.04 ± 0.03 | 0.04 ± 0.02 | 0.84 ± 0.06 | 0.01 ± 0.01 | 0.01 ± 0.01 | 0.01 ± 0.01 | 0.01 ± 0.01 | 0.07 ± 0.02 | 0.85 ± 0.04 | 0.03 ± 0.01 | 0.01 ± 0.01 |
| | | 5mer | 0.02 ± 0.04 | 0.04 ± 0.04 | 0.04 ± 0.03 | 0.01 ± 0.01 | 0.82 ± 0.07 | 0.01 ± 0.00 | 0.01 ± 0.03 | 0.02 ± 0.03 | 0.02 ± 0.02 | 0.04 ± 0.02 | 0.86 ± 0.07 | 0.01 ± 0.01 |
| | | 6mer | 0.02 ± 0.04 | 0.03 ± 0.04 | 0.03 ± 0.03 | 0.03 ± 0.02 | 0.01 ± 0.00 | 0.81 ± 0.09 | 0.03 ± 0.04 | 0.03 ± 0.04 | 0.02 ± 0.02 | 0.03 ± 0.02 | 0.02 ± 0.01 | 0.82 ± 0.10 |

**Table S3 Reaction rate constants for EGFR pre-dimerization.**

| | | | -EGF | | | | | | | | | |
|---|---|---|---|---|---|---|---|---|---|---|---|---|
| | | | | | | | | | # of cells : n= 13 | | | |
| | | | immobile | | | slow-mobile | | | | fast-mobile | | |
| control | Dissociation rate constant | dimerization | 1.1 | ± | 0.8 | 0.7 | ± | 0.2 | | 8.9 | ± | 2.8 |
| | | decomposition | 4.4 | ± | 0.6 | 7.6 | ± | 0.5 | | 11.3 | ± | 2.8 |
| | | | | | | | | | | n= 6 | | |
| MβCD 10 mM | | | immobile | | | slow-mobile | | | | fast-mobile | | |
| | Dissociation rate constant | dimerization | 1.2 | ± | 0.5 | 0.6 | ± | 0.1 | | 14.4 | ± | 4.8 |
| | | decomposition | 4.5 | ± | 0.5 | 6.1 | ± | 0.5 | *** | 14.3 | ± | 3.7 |
